# Human stem cell model of *HNF1A* deficiency shows uncoupled insulin to C-peptide secretion with accumulation of abnormal insulin granules

**DOI:** 10.1101/2021.01.26.428260

**Authors:** Bryan J. González, Haoquan Zhao, Jacqueline Niu, Damian J. Williams, Jaeyop Lee, Chris N. Goulbourne, Yuan Xing, Yong Wang, Jose Oberholzer, Xiaojuan Chen, Charles A. LeDuc, Wendy K. Chung, Henry M. Colecraft, Jesper Gromada, Yufeng Shen, Robin S. Goland, Rudolph L. Leibel, Dieter Egli

## Abstract

Mutations in *HNF1A* cause Maturity Onset Diabetes of the Young type 3 (MODY3), the most prevalent form of monogenic diabetes. We generated stem cell-derived pancreatic endocrine cells from human embryonic stem cells (hESCs) with induced hypomorphic mutations in *HNF1A*. Using these cells, we show that *HNF1A* orchestrates a transcriptional program required for distinct aspects of β-cell fate and function. During islet cell differentiation, *HNF1A* deficiency biases islet endocrine cells towards an *α*-cell fate associated with *PAX4* down-regulation. *HNF1A-* deficient β-cells display impaired basal and glucose stimulated-insulin secretion in association with a reduction in *CACNA1A* and intracellular calcium levels, and impaired insulin granule exocytosis in association with *SYT13* down-regulation. Knockout of *PAX4*, *CACNA1A* and *SYT13* reproduce the relevant phenotypes. Reduction of insulin secretion is associated with accumulation of enlarged secretory granules, and altered stoichiometry of secreted insulin to C-peptide. In *HNF1A* deficient β-cells, glibenclamide, a sulfonylurea drug used in the treatment of MODY3 patients, increases intracellular calcium to levels beyond those achieved by glucose, and restores C-peptide and insulin secretion to a normal stoichiometric ratio. To study *HNF1A* deficiency in the context of a human disease model, we also generated stem cell-derived pancreatic endocrine cells from two MODY3 patient’s induced pluripotent stem cells (iPSCs). While insulin secretion defects are constitutive in cells with complete *HNF1A* loss of function, β-cells heterozygous for hypomorphic *HNF1A* mutations are initially normal, but lose the ability to secrete insulin and acquire abnormal stoichiometric secretion ratios. Importantly, the defects observed in these stem cell models are also seen in circulating proportions of insulin:C-peptide in nine MODY3 patients.

**One sentence of summary:** Deficiency of the transcription factor *HNF1A* biases islet endocrine cell fate towards *α*-cells, impairs intracellular calcium homeostasis and insulin exocytosis, alters the stoichiometry of insulin to C-peptide release, and leads to an accumulation of abnormal insulin secretory granules in β-cells.

## Introduction

Maturity onset diabetes of the young (MODY) is an autosomal dominant form of diabetes with onset typically before the age of 25 years accounting for 1-2% of diabetes incidence (*1*). There are at least 11 (*2*) genetically distinct types of MODY, due to derangements in β-cell development or function. MODY3 is caused by mutations in the transcription factor HNF1A (*3, 4*) and accounts for 63% of diagnosed instances of MODY (*5*). MODY3 patients have normal glucose tolerance during childhood and early adult life, but show progressive reduction of insulin secretion in response to glucose (*6–9*). Glycemia typically increases over time, resulting in the need for treatment with the insulin secretory sulfonylurea drugs. Eventually, 30-40% of patients become insulin-dependent due to progressive deterioration of β-cell function.

HNF1A is a 631-amino acid transcription factor with three major domains: dimerization, DNA-binding, and transactivation. Over 200 *HNF1A* diabetes-related mutations have been identified in all major ethnic groups (*10*) and the DNA-binding domain has the highest frequency of mutation in MODY3 patients (*5*). Understanding the role of *HNF1A* gene and the pathophysiology of MODY3 has been difficult because of the difficulty in studying human islets or mass with these mutations. Mouse models of HNF1A deficiency do not accurately mimic patient phenotypes (*11*).

Stem cell-derived β-cells provide a useful model system, and have been used to study β-cell development in humans (*12, 13*) and to recapitulate disease phenotypes (*14, 15*). Differentiation of pluripotent stem cells to pancreatic endocrine cells can be achieved by a multistep protocol resulting in islet-like clusters containing all endocrine cell types (*16–18*). Transplanting these islet-like clusters into mice allows functional testing of stem cell-derived β-cells (scβ-cells) *in vivo* (*19, 20*). Here we show that human stem cell-based models of *HNF1A* deficiency shows islet developmental bias towards *α*-cells, altered insulin granule morphology and the stoichiometry of insulin:C-peptide secretion *in vitro* and *in vivo*. We used these models to identify disordered calcium homeostasis and glucose-mediated exocytosis as key mechanisms accounting for the secretory defects observed. This study was designed using two cell-based models: Human embryonic stem cell (hESC) lines null for *HNF1A* in **Fig. 1-5** and **Fig. S1**-**S10**; and induced pluripotent stem cell (iPSC) lines with MODY3 patient-specific mutations in **Fig. 6** and **Fig. S11**-**S13**. Measurements of circulating hormones in nine MODY3 patients are presented in **Fig. 7**. **Fig. S1** provides a schematic overview of the study.

**Fig. 1.**
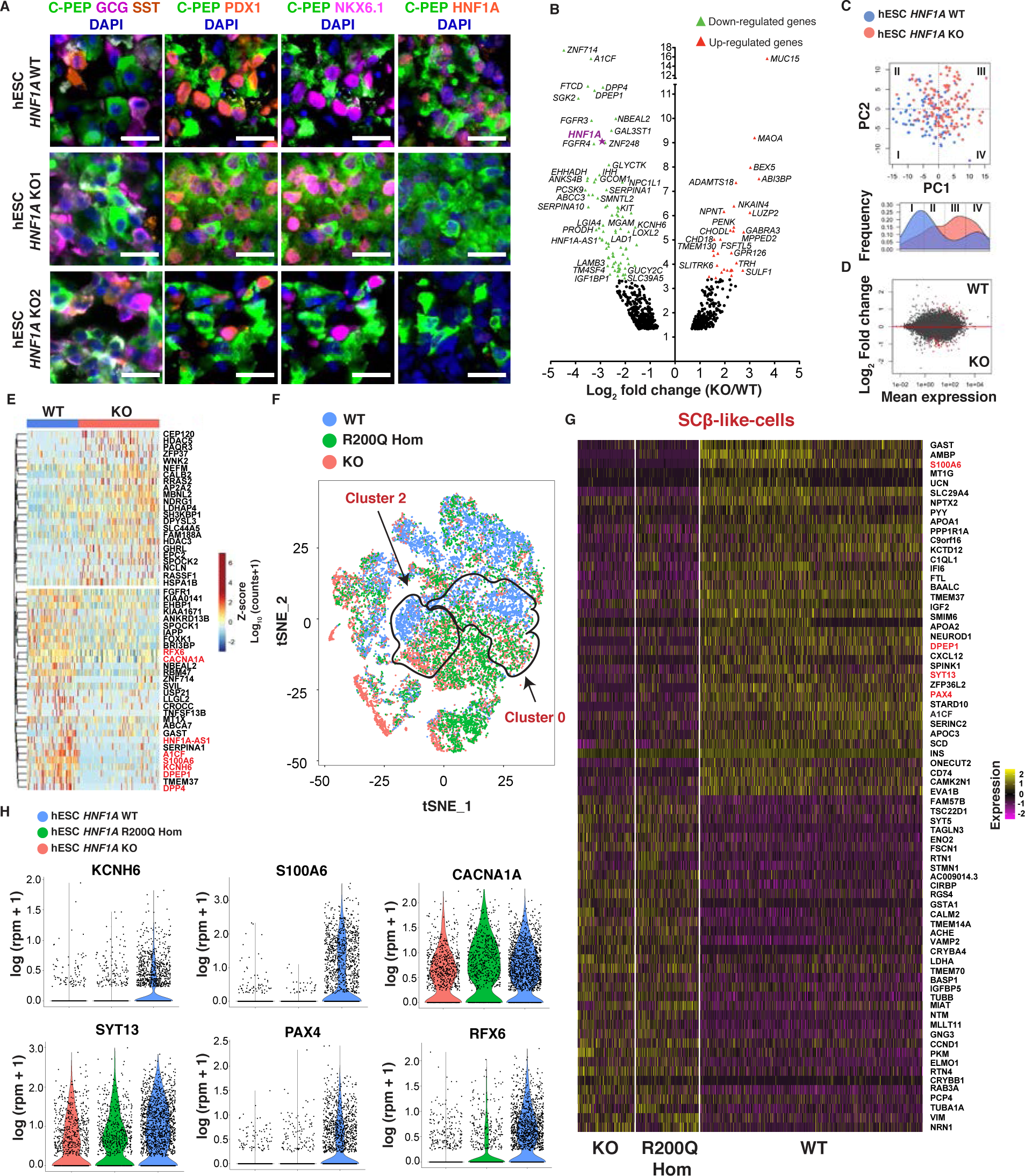
*HNF1A* deficiency impairs a network of genes required for calcium signaling, glucosestimulated insulin exocytosis and β-cell fate *in vitro*. (A) Representative immunohistochemistry (IHC) images of hESC-derived endocrine cell lines (*HNF1A* WT, KO1 and KO2) for indicated markers. White cells are GCG/CPEP double positive cells. Scale bars: 20 µm. **(B-E)** RNA sequencing of FACs sorted *INS^GFP/wt^* positive cells between *HNF1A* WT and KO genotypes (n=3 for both genotypes). **(B)** Volcano plot depicting fold change (log_2_ fold change, x-axis) and statistical significance (-log_10_ p-value, y-axis) for differential gene expression (down-regulated in green; up-regulated in red) by bulk RNAseq (see also Table S3). **(C)** Cell clustering by principal component analysis (PCA). Lower panel depicts the radial distribution or frequency of individual cells from each PCA quadrant. **(D)** MA-plot depicting fold change (log_2_ fold change, y-axis) and mean expression (counts, x-axis) of differentially-expressed genes. **(E)** Heatmap showing expression for each gene identified as z-score of expression from all sorted scβ-like cells. **(F)** Single cell RNA sequencing of 22164 (all genotypes combined) unsorted hESC-derived endocrine cells by *HNF1A* WT, R200Q homozygous and KO genotypes (n=3 for each genotype). Feature plot based on tSNE projection of cells where the colors denote different cell lines by *HNF1A* genotype line via Louvain algorithm performed by Seurat. **(G)** Heatmap showing differentially expressed genes from scβ-like cells by *HNF1A* WT, R200Q homozygous and KO genotypes. Total of 1846 scβ-like cells (all genotypes combined) were identified and displayed as |logFC|>0.35 and adjusted p-value <1e^-4^. Genes are listed in decreasing order of log_2_ fold change between *HNF1A* WT and *HNF1A* mutant genotypes. **(H)** Violin plot of cells based on *KCNH6*, *S100A6*, *CACNA1A*, *SYT13*, *PAX4* and *RFX6* gene expression (log1(rpm+1)) from scβ-like cells. All stem cell differentiations were done for 27 days. (n) represents the number of biological replicates. See also Fig. S4 and S5.

## Results

### Generation of isogenic cell lines with *HNF1A* mutations in hESCs and MODY3 iPSCs

To elucidate the cellular functions of *HNF1A*, we used CRISPR/Cas9 technology to generate hESC lines (Mel1) harboring non-naturally occurring heterozygous and homozygous null mutations. The Mel1 hESC line has a *INS^GFP/wt^* and *GAPDH^Luciferase/wt^* dual-reporter containing a GFP gene knockin in place of a coding *INS* locus, and a luciferase knockin in one *GAPDH* locus (*21*). GFP expression under the control of the insulin promoter does not alter *β*-cell function in mice (*22*) and does not cause ER-stress in human sc*β*-like-cells (*23*). This marker enables imaging and isolation of viable INS-GFP^+^ cells *in vitro* and *in vivo*. We designed short guide RNAs (sgRNAs) to introduce indel mutations in the DNA-binding domain (exon 3) of the *HNF1A* gene (**Fig. S2A** and **Table S1**) because the DNA-binding domain has the highest frequency of mutation in MODY3 patients (*5*). Transfection of hESC lines with Cas9-GFP and sgRNAs #12 or #14, followed by sorting for GFP (**Fig. S2B**), achieved 17.3 and 20.4% indel efficiency for the sgRNAs as shown by surveyor assay (**Fig. S2C**). Gene editing was efficient, resulting in compound heterozygous knockouts and heterozygous mutations (**Fig. S2D**) with no off-target mutations detected through targeted sequencing (**Table S2**). Heterozygous (hESC *HNF1A* Het) and compound heterozygous-null mutant cell lines (hESC *HNF1A* KO1 and KO2) with different indels in exon 3 and premature protein termination were chosen for further studies (**Fig. S2E**). hESC *HNF1A* Het, KO1 and KO2 lines retained the HNF1A dimerization domain (truncated HNF1A), while hESC *HNF1A* KO3 line was mutated at the start codon, deleting the entire HNF1A protein (**Fig. S2E**).

To understand the consequences of MODY3 patient-specific heterozygous mutations, we also generated iPSC lines from MODY3 patient fibroblasts containing heterozygous mutations in the transactivation domain (MODY3 iPSC Het: +/460_461insCGGCATCCAGCACCTGC); and DNA-binding domain (MODY3 iPSC Het: +/R200Q). The R200Q variant has been previously associated with MODY3 (*24*); however, the precise effect of this missense mutation on *HNF1A* function is unknown, but very likely pathogenic (*24*). Using CRISPR/Cas9, we corrected the R200Q mutation from MODY3 iPSC Het (**Fig. S2F**) with an efficiency of 7.9% (5 clones out of 63) to generate isogenic wildtype cells. In addition to the hESC knockout lines, we also generated iPSCs compound heterozygous knockout lines with an efficiency of 42.9% (27 clones out of 63) (**Fig. S2D**). An iPSC line with a deletion of the WT allele, but intact patient’s mutation (MODY3 iPSC: R200Q/-) was chosen for further studies (**Fig. S2G**). We also introduced the same R200Q mutation into both alleles in the hESC line (hESC Hom: R200Q/R200Q) (**Fig. S2H**) with an efficiency of 20.8% (5 clones out of 24) (**Fig. S2I**). All genetically manipulated cell lines (**Fig. S2J**) resulted in modified versions of HNF1A protein (**Fig. S2K and S2L**) and had a normal karyotype (**Fig. S2M and S2N**) and no off-target mutations (**Table S2**).

### *HNF1A* is not required to generate pancreatic endocrine cells

To determine the functional consequences of *HNF1A* deficiency, the first part of this study will focus on the hESC model. Stem cells were differentiated into the pancreatic lineage using a published differentiation protocol (*20*) (**Fig. S3A**). Differentiation of wildtype stem cells using this protocol consistently generated 82% PDX1^+^/NKX6.1^+^ cells at the pancreatic progenitor stage (day 11). At the endocrine stage (day 27), we obtained 45% PDX1^+^/CPEP^+^ cells with 60% of them co-expressing NKX6.1 (sc*β*-like cells), 30% GCG^+^ cells (sc*α*-like cells) and 10% SST^+^ cells (scδ-like cells) (**Fig. S3B**).

To determine the timing of *HNF1A* expression, we performed qPCR at different stages of differentiation. Insulin mRNA was detected at the endocrine stage (day 20) (**Fig. S3C**), whereas *HNF1A* mRNA is first detected at low levels at the primitive gut tube stage (day 5), increased at the pancreatic progenitor stage (day 11) and increased further at the endocrine stage (day 27) (**Fig. S3D**), identifying the stages where lack of *HNF1A* could have developmental and/or functional consequences. *HNF1A* knockout resulted in significant reduction of *HNF1A* transcript at the endocrine stage (**Fig. S3E**); HNF1A protein was detected in hESC WT, but was absent in hESC *HNF1A* KO-derived endocrine cells by western blot (**Fig. S3F**).

Mutations in *HNF1A* did not affect hESC lines in generating definitive endoderm cells (SOX17^+^) at day 3 of differentiation or pancreatic progenitor cells (PDX1^+^/NKX6.1^+^) at day 11 of differentiation as determined by immunohistochemistry (**Fig. S3G**). At the endocrine stage of development (day 27), cell clusters or organoids were morphologically indistinguishable. hESC lines became positive for INS-GFP fluorescence due to the *INS^GFP/wt^* reporter; there was no differences in GFP expression by genotype (**Fig. S3H**). *HNF1A* KO cells were capable of differentiation to all islet endocrine cell types, including PDX1^+^/NKX6.1^+^/SYP^+^/CPEP^+^ cells (sc*β*-like cells), GCG^+^ cells (sc*α*-like cells) and SST^+^ cells (scδ-like cells) (**Fig. 1A and S3I-S3K**) indicating that *HNF1A* is not required to generate pancreatic endocrine cells. No differences were found for PDX1 and NKX6.1 in hESC *HNF1A* WT and *HNF1A* KO endocrine cells (day 27) by immunohistochemistry (**Fig. S3J and S3K**) and flow cytometry (**Fig. S3L**). The ability of *HNF1A* KO cells to differentiate to all endocrine cell types enabled the comparison of transcriptional programs in *HNF1A* KO and WT cells.

### *HNF1A* deficiency impairs a network of genes required for calcium signaling, glucose-stimulated insulin exocytosis and β-cell fate

To determine the transcriptional consequences of *HNF1A* deficiency in scβ-cells, bulk and single-cell RNA sequencing was performed in INS-GFP sorted β-like cells derived from hESC *HNF1A* KO and WT lines *in vitro*. Volcano plot analysis of bulk RNA sequencing identified 30 up-regulated genes and 148 down-regulated genes, including *HNF1A* in hESC *HNF1A* KO β-like cells (**Fig. 1B and Table S3**). Single-cell RNA sequencing showed that hESC *HNF1A* KO β-like cells segregated from hESC *HNF1A* WT β-like cells based on their gene expression profile by principal component analysis (**Fig. 1C**), and volcano plot analysis (**Fig. 1D**) revealed previously undescribed HNF1A target genes involved in intracellular calcium signaling (*CACNA1A*, *S100A6* and *TMEM37*), calcium-mediated insulin secretion (*KCNH6*) and glucose-stimulated insulin secretion (*RFX6*, *DPEP1*, *DPP4* and *IAPP*) (**Fig. 1E**) as commonly downregulated with bulk RNA sequencing from *HNF1A* KO scβ-like cells. Among the up-regulated genes in hESC *HNF1A* KO β-like cells, we found genes involved in synaptic vesicle cycle (*AP2A2*, *SH3KBP1* and *HSPA1B*) (**Fig. 1E and Table S4**).

To understand the transcriptional consequences of *HNF1A* mutation in other endocrine cell types, we analyzed all single cells within the islet-like clusters, allowing the segregation of cell types according to single-cell gene expression. In addition to *HNF1A* KO lines, an hESC line homozygous for *HNF1A* R200Q (R200Q/R200Q), a MODY3 patient-specific mutation, was included. Using clustering analysis from those cells, we grouped the cells into thirteen different cell type populations (**Fig. S4A and S4B**). To cluster different endocrine cells, we used the expression of endocrine cell markers such as *SYP*, *INS*, *GCG* and *SST* (**Fig. S4C**) and pancreatic progenitor markers *NKX6.1*, *PDX1*, *MAFA* and *HNF1A* (**Fig. S4D**) from all cell lines. This allowed the identification of insulin expressing cells (cluster 0) and glucagon expressing cells (cluster 2) (**Fig. 1F**). From the population of insulin expressing cells (cluster 0), cells defined by high insulin and low glucagon expression were considered scβ-like cells and were compared between genotypes (**Fig. S5A**).

Separation into populations according to these markers enabled us to identify differentially expressed genes in *HNF1A* mutant cells affecting specific endocrine cell types. In scβ-like cells, a total of 73 genes were differentially expressed between *HNF1A* WT, homozygous R200Q and KO lines (**Fig. 1G and Table S5**). Pathway analysis of differentially expressed genes in scβ-like cells revealed genes involved in MODY, endocrine cell development, calcium signaling/sensing, insulin secretion and synaptic vesicle cycle (**Fig. S5B**). Furthermore, several MODY genes (*RFX6*, *PAX4* and *NEUROD1*) were down-regulated (**Fig. 1G**). These transcription factors are known to be important for the functional identity of adult pancreatic β-cells (*25, 26*). We also identified down-regulated genes important for intracellular calcium signaling (*S100A6* and *TMEM37*), exocytosis-regulated insulin secretion (*SYT13*) and glucose-stimulated insulin secretion (*IGF2* and *LLGL2*) in hESC *HNF1A* mutated β-like cells (**Fig. 1I and Table S5**) (*25, 27, 28*). Among the up-regulated genes, we found genes involved in synaptic vesicle cycle (*RAB3A* and *VAMP2*) (**Fig. 1G and Table S5**). *RFX6*, *KCNH6*, *S100A*6, *A1CF*, *DPEP1* and *DPP4* were the most consistently downregulated genes in hESC *HNF1A* mutated β-like cells. In addition to those genes, *SYT13*, *CACNA1A* and *PAX4* were down-regulated in the scβ-like-cell subpopulation (**Fig. 1H**). Similar to our study, a recent case study of cadaveric human islets of a MODY3 patient (+/T260M) also found down-regulation of *RFX6*, *CACNA1H*, *IAPP* and *TMEM37* in β-cells, and down-regulation of *CACNA1A*, *RFX6*, *PCBD1* and *PPP1R1A* in *α*-cells (*29*). These genes are required for calcium signaling, glucose-stimulated insulin exocytosis, and endocrine cell fate.

To determine the molecular consequences of *HNF1A* deficiency in other endocrine cell types, we analyzed mono-hormonal sc*α*-like cells (cluster 2a), characterized by high glucagon and low insulin expression (**Fig. S5C and S5D**). We identified up-regulated genes involved in glucagon signaling pathways (*PKM*, *CALM1* and *CALM2*) and down-regulated genes involved in insulin secretion (*ADCY1*, *KCNH6*, *RFX6*, *IGF2*, *TM4SF4* and *ATF4*), calcium-mediated exocytosis (*SYT7*) and β-cell dedifferentiation (*GC*) (*30*) (**Fig. S5E and S5F**). Furthermore, in bi-hormonal cells expressing both insulin and glucagon, *PYY* was down-regulated, while *GCG* was up-regulated (**Fig. S5G and S5H**) in *HNF1A* mutated lines, indicating a transcriptional bias of endocrine cells towards the *α*-cell fate.

A recent publication by Cardenas-Diaz et al. (*31*) identifies *LINC01139* as an *HNF1A* target implicated in β-cell dedifferentiation and β-cell respiration. However, we found no significant difference in the expression of this long non-coding RNA between WT and *HNF1A* mutated (hESC Het, hESC Hom R200Q/R200Q, hESC KO or iPSC MODY3 Het +/460ins) β-like (**Fig. S5I-5K**) and *α*-like cells (hESC Hom R200Q/R200Q and hESC KO) (**Fig. S5L**). Similar results were reported in cadaveric human β-cells from non-diabetic donors as compared to one MODY3 donor (+/T260M) (**Fig. S5M**).

In summary, *HNF1A* orchestrates a network of genes promoting endocrine differentiation to β-cells, and in differentiated β-cells regulates genes involved in calcium signaling, hormone exocytosis and glucose-stimulated insulin secretion.

### *HNF1A* deficiency causes a developmental bias towards the *α*-cell fate

Downregulation of genes involved in adult β-cell identity (*PAX4* and *RFX6*) with transcriptional bias towards the *α*-cell fate in *HNF1A* KO cells point to a developmental role of HNF1A. Though both hESC *HNF1A* KO and WT cells gave rise to similar numbers of insulin positive cells (**Fig. S6A and S6B**), immunostaining showed a variable increase in sc*α*-like cell numbers from 30% to up to 70% in the hESC *HNF1A* KO line (**Fig. 2A and S6B**). This was confirmed by a significant increase in GCG^+^/CPEP^+^ cell type ratio by immunohistochemistry (**Fig. 2B**) and glucagon content by ELISA (**Fig. 2C**). Furthermore, quantification by flow cytometry consistently showed a significant increase in sc*α*-like cells in three different hESC *HNF1A* KO lines as compared to hESC *HNF1A* WT lines (**Fig. S6C and S6D**). The absolute increase in sc*α*-like cells in hESC *HNF1A* KO cells was driven by an increase in the number of bi-hormonal (56% in KO vs. 20% in WT) and mono-hormonal (7% vs. 1%) sc*α*-like cells as shown by flow cytometry (**Fig. 2D**), immunohistochemistry (**Fig. S6E**), and RNA sequencing (**Fig. S6F**). Transcriptional analysis by single-cell RNA sequencing of glucagon and insulin double hormone-positive cells showed an increase in glucagon gene expression and a decrease in insulin gene expression in hESC *HNF1A* KO cells compared to WT cells (**Fig. 2E**). These findings are consistent with *PAX4* repressing pancreatic glucagon gene expression (*32*), and indicate that *HNF1A* hypomorphism affects endocrine cell fate by increasing expression of the glucagon gene in pancreatic endocrine cells. *HNF1A* deficiency did not affect the proportion of scδ-like cell fate as shown by flow cytometry (**Fig. S6G and S6H**). No effects of *HNF1A* mutations on proliferation (**Fig. S6I**) or survival (**Fig. S6J and S6K**) of sc*β*-like cells were observed by flow cytometry. These results show that *HNF1A* deficiency biases endocrine cells toward the *α*-cell fate *in vitro*.

**Fig. 2.**
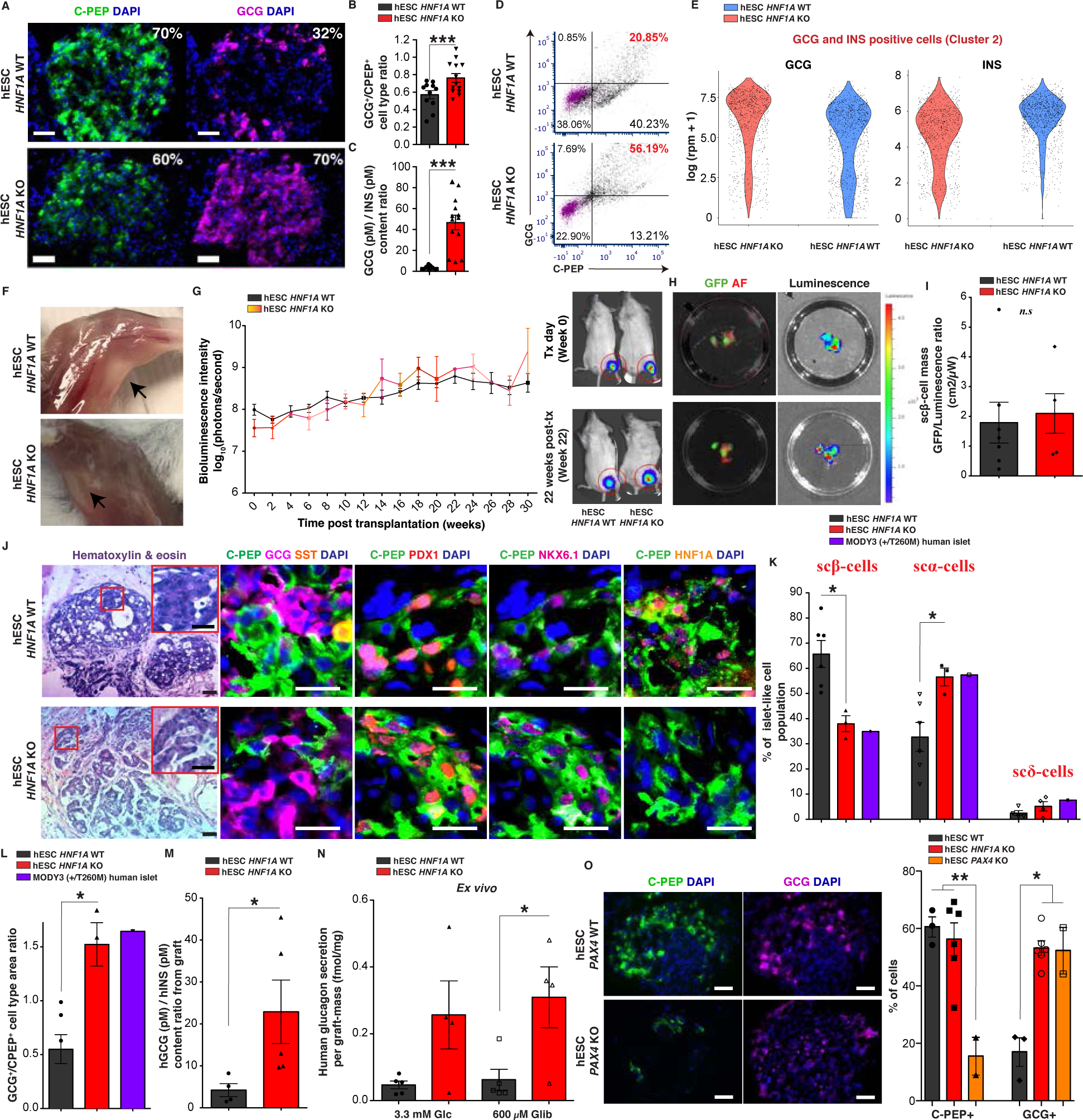
*HNF1A* deficiency causes a developmental bias toward the α-cell fate. **(A)** Representative IHC images of hESC-derived endocrine cell lines (*HNF1A* WT and KO) for indicated markers. Scale bars: 50 μm. **(B)** GCG^+^/CPEP^+^ cell type ratios by IHC and **(C)** GCG/INS protein content ratios measured by ELISA. p-values: ***p<0.001; two-tailed t-test. (**D)** CPEP^+^ and GCG ^+^ populations by flow cytometry. Gating for GCG and CPEP negative cells (magenta) was determined by incubating cells without primary antibodies and with secondary antibodies. **(E)** Violin plot of total of 2670 (all genotypes) GCG and INS positive cells (cluster 2) displayed based on *GCG* and *INS* expression (log(rpm+1)) (n=3 for each genotype). **(F)** Representative image of graft tissue (black arrows) 30 weeks post-transplantation with hESC-derived endocrine cells before explant from quadriceps muscle. **(G)** Bioluminescence intensity (log_10_ photons/second) measured in mice transplanted with hESC-derived endocrine cells (WT n=11 and KO n=4) over time (weeks) with representative bioluminescence images. **(H)** Representative GFP fluorescence (scβ-cells) and bioluminescence images (total cells) (GFP in green; AF: tissue auto-fluorescence in red) from isolated grafts with **(I)** quantification of scβ-cell mass GFP/luminescence ratio (cm^2^/µW). **(J)** Representative H&E and IHC images showing hESC-derived endocrine cells for indicated markers. Scale bars: 20 µm. **(K)** Percentage of scβ-, sc*α*- and scδ-cells from the islet-like clusters and MODY3 islets (*HNF1A* +/T260M) from Haliyur et al. (*29*). **(L)** GCG^+^/CPEP^+^ cell type area ratios by IHC from isolated grafts and GCG^+^/INS^+^ from MODY3 islets (*HNF1A* +/T260M, Haliyur et al). Each point in plots is an islet-like cluster. Endocrine cell percentages were calculated as hormone-positive area from islet-like clusters across the entire graft (2 grafts for each genotype). **(M)** hGCG (pM)/hINS (pM) content ratio from isolated grafts. **(N)** *Ex vivo* human glucagon secretion normalized to graft mass (fmol/mg) in response to indicated secretagogues. All protein concentrations were measured by ELISA. **(O)** Representative IHC images of hESC-derived endocrine cell lines for indicated markers with quantification of CPEP+ and GCG+ cells (%) by flow cytometry. All stem cell differentiations were done for 27-30 days. 20 clusters (∼10k cells per cluster) of endocrine cells were used flow cytometry. All mice were transplanted with clusters of hESC-derived endocrine cells (*HNF1A* WT and KO2) at day 27 of differentiation, and grafts were isolated 30 weeks post-transplantation for *ex vivo* analysis. (n) represents the number of biological replicates. For scatter plots, each point in plots represents an independent biological experiment (n). Data are represented as mean ± SEM. p-values: *p<0.05, **p<0.01, ***p<0.001; Mann-Whitney test. n.s: non-significant. See also Fig. S6 and S7.

In order to assess the developmental requirements of *HNF1A* in endocrine cells *in vivo*, we transplanted pancreatic islet-like clusters derived from hESC *HNF1A* WT and hESC *HNF1A* KO lines with *GAPDH^Luciferase/wt^* and *INS^GFP/wt^* dual-reporter (*21*). Mice received ∼180 clusters of stem cell-derived islet-like cells (**Fig. S7A**) containing similar amount of CPEP/PDX1 positive cells across genotypes (**Fig. S7B and S7C**); but with different amounts of bihormonal GCG/CPEP positive cells (**Fig. 2D**). Cell Transplantation was done by injection into the ventral and medial muscle groups of the left quadriceps in NOD SCID gamma immunodeficient mice (NSG mice) (**Fig. 2F**). The skeletal muscle was chosen for its dense vasculature and easy access for surgical procedures. Skeletal muscle has been used for other endocrine transplants, including for parathyroid auto-transplantation in patients undergoing parathyroidectomy with a 93% success rate (*33*). Analysis of engraftment efficiency was evaluated by bioluminescence intensity (BLI) several weeks post-transplantation. Mice with successful engraftment showed a two-fold increase in BLI 4 to 6 weeks post transplantation, while those with failed engraftment showed a decrease (**Fig. S7D**). Transplantation was successful in 79% (31/39) of mice, independent of *HNF1A* genotype (**Fig. S7E**). 92.5% (49/53) of mice transplanted with hESC lines remained teratoma-free (**Fig. S7F**). In those mice, the graft explant (30 weeks post-transplantation) (**Fig. S7G**) was a vascularized tissue of about ∼220 mg (**Fig. S7H**). According to luminescence intensity, there was a gradual increase in BLI from week 0 to 30 post-transplantation without significant differences by *HNF1A* genotype (**Fig. 2G**). Grafts remained localized for up to 50 weeks post-transplantation, and in no case (0/39) was luminescence detected in an ectopic location. These results show that skeletal muscle is a stable and suitable transplantation site for SC-derived islet-like cells.

Thirty weeks post-transplantation, we isolated grafts from normoglycemic animals. Quantification of sc*β*-mass as determined by GFP fluorescence did not differ by *HNF1A* genotype (**Fig. 2H-2I and S7I-S7J**). The presence of exclusively mono-hormonal endocrine cells, including sc*β*-cells (CPEP^+^/PDX1^+^/NKX6.1^+^), as well as glucagon- and somatostatin-positive cells were apparent in both hESC *HNF1A* WT and hESC *HNF1A* KO grafts (**Fig. 2J**). No double hormone-positive cells were identified in cells of either genotype. Consistent with the *in vitro* studies, the percentage of sc*α*-cells in *HNF1A* KO compared to *HNF1A* WT islet-like structures were increased by 24% (**Fig. 2K and S7K**), leading to a significant increase in GCG^+^/CPEP^+^ cell type ratio (**Fig. 2L**); similar results as found in cadaveric human islets of a MODY3 donor (+/T260M) (*29*) (**Fig. 2K**). Consistent with this observation, *ex vivo* analysis of isolated hESC *HNF1A* KO grafts showed an increase in glucagon protein content by ELISA (**Fig. 2M and S7L-S7M**). Glucagon secretion was higher in hESC *HNF1A* KO grafts when stimulated with glibenclamide, a second-generation sulfonylurea drug compared to hESC *HNF1A* WT grafts (**Fig. 2N**). Similarly, elevated glucagon secretion upon KCl stimulation was detected *in vitro* from hESC *HNF1A* KO-derived endocrine cells (**Fig. S7N).** These results point not only to a gain of *α*-cell number, but also enhanced *α*-cell function due to *HNF1A* deficiency. This is consistent with the up-regulation of genes (*PKM*, *CALM1* and *CALM2*) involved in glucagon signaling pathways in sc*α*-like cells.

To determine whether the downregulation of *PAX4* seen in *HNF1A* mutant cells (**Fig. 1H**) could reproduce the bias towards the *α*-cells, we generated hESC *PAX4* KO cell lines using CRISPR/Cas9 (**Fig. S7O**). The *PAX4* gene is an essential gene for differentiation of insulin-producing *β*-cells in the mammalian pancreas (*34*). Knockout of *PAX4* in rabbits induces a decreased in number of *β*-cells and increased number of *α*-cells (*26*), whereas overexpression of *PAX4* reduces glucagon expression in differentiating hESCs (*35*). Endocrine cells differentiated from an hESC *PAX4* KO line were characterized by significant decrease in sc*β*-like cells compared to hESC WT and *HNF1A* KO lines. In the hESC *PAX4* KO lines, sc*α*-like cells were increased to levels similar to hESC *HNF1A* KO cells by immunohistochemistry (**Fig. 2O**) and flow cytometry (**Fig. S7P**). Therefore, in *HNF1A* KO lines, down-regulation of *PAX4* and de-repression of glucagon gene are associated with the developmental bias towards *α*-cell fate.

In summary, *HNF1A* is not required for development of mono-hormonal endocrine cells (*α*-, *β* and δ-cells). But restriction of HNF1A activity increases the number and proportion of *α*-cells in association with *PAX4* downregulation. As *HNF1A* clearly regulates a network of genes, a role of additional *HNF1A* target genes in *β*-and *α*-cell fate is likely.

### *HNF1A* deficiency affects glucose-mediated exocytosis of insulin granules

To determine the consequences of *HNF1A* deficiency on β-cell function, we measured glucose-stimulated insulin secretion. hESC *HNF1A* KO sc*β*-like cells had reduced basal insulin secretion (**Fig. S8A**) and impaired glucose-stimulated insulin secretion compared to WT sc*β*-like cells and compared to human pancreatic islets *in vitro* (**Fig. 3A**). These differences in hormone secretion between *HNF1A* mutant and *HNF1A* WT scβ-like cells were not due to a reduction in insulin content (**Fig. S8B**). In contrast to glucose, treatment with the insulin secretagogues, tolbutamide or potassium chloride resulted in insulin secretion in hESC *HNF1A* KO cells comparable to hESC *HNF1A* WT *β*-like cells (**Fig. 3B**). Thus, membrane depolarization enables recruitment of an insulin granule reservoir that is abnormally retained in *HNF1A* mutant *β*-cells and fails to be secreted in response to glucose. The ability of sulfonylureas in releasing insulin from β-cell granules is consistent with the clinical efficacy of sulfonylurea drugs in MODY3. These defects are likely intrinsic to the *β*-cell, and not due to paracrine effects of glucagon (*36*).

**Fig. 3.**
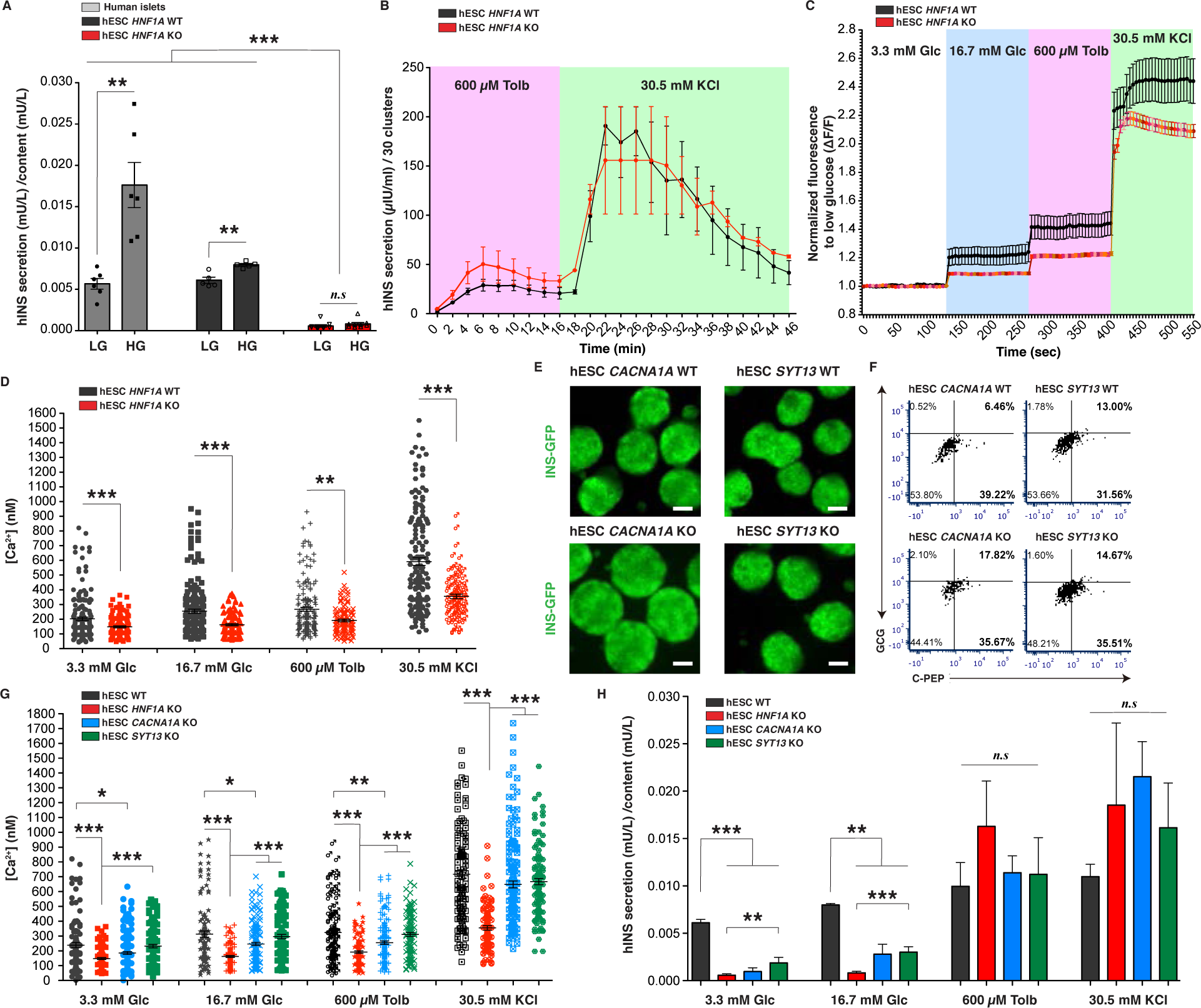
*HNF1A* deficiency affects insulin secretion due to reduced intracellular calcium levels in association with *CACNA1A* and *SYT13* down-regulation *in vitro*. (A) Human insulin secretion (mU/L) in 1h normalized to content (mU/L) in response to low glucose (LG, 3.3 mM) and high glucose (HG, 16.7 mM) stimulation in static assay of hESC-derived endocrine cell lines (*HNF1A* WT and KO) and pancreatic human islets. **(B)** Human insulin secretion (μIU/ml) normalized to 30 clusters in a perfusion assay (both *HNF1A* WT and KO genotypes n=2) in response to indicated secretagogues. **(C)** Intracellular calcium fluorescence normalized to low glucose (F340/380) from dispersed scβ-like cells with **(D)** absolute intracellular calcium concentration (nM) quantification in response to indicated secretagogues (each point represents a scβ-like-cell, *HNF1A* WT n=170 and *HNF1A* KO n=110 cells). **(E)** Representative GFP-field images of endocrine organoids from indicated genotypes. Scale bars: 200 µm. **(F)** Representative CPEP^+^ and GCG ^+^ populations quantification by flow cytometry. **(G)** Absolute intracellular calcium concentration (nM) in response to indicated secretagogues (each point represents a scβ-like-cell from independent batches of differentiation, WT n=110, *HNF1A* KO n=110, *CACNA1A* KO n=175 and *SYT13* KO n=120 cells). **(H)** Human insulin secretion (mU/L) in 1h normalized to content (mU/L) in response to low (3.3 mM) and high glucose (16.7 mM) and indicated secretagogues in static assay (WT n=5, *HNF1A* KO n=9, *CACNA1A* KO n=6 and *SYT13* KO n=5). All stem cell differentiations were done for 27 days. 20 clusters (∼10k cells per cluster) of endocrine cells were used flow cytometry. (n) represents the number of biological replicates. For scatter plots, each point in plots represents an independent biological experiment (n). Data are represented as mean ± SEM. p-values: *p<0.05, **p<0.01, ***p<0.001; Mann-Whitney test. n.s: non-significant. See also Fig. S8.

As calcium signaling is a mechanism for the release of insulin through granule exocytosis, we measured intracellular calcium levels in dispersed sc*β*-like cells *in vitro*. Both genotypes had glucose-stimulated calcium responses (**Fig. 3C**). However, hESC *HNF1A* KO *β*-like cells had significantly lower calcium levels relative to hESC *HNF1A* WT sc*β*-like cells as measured by Fura-2 fluorescence (**Fig. 3C**). This was further supported by measurements of absolute intracellular calcium concentrations by Fura-2 fluorescence (**Fig. 3D**). Among the HNF1A target genes, reduced expression of *CACNA1A* is potentially involved in reducing intracellular calcium levels, and *SYT13* may be required for efficient exocytosis-mediated insulin secretion. *CACNA1A* encodes a voltage-dependent calcium channel mediating the entry of calcium into excitable cells and is involved in calcium-dependent insulin secretion and type 2 diabetes (*37*). Synaptotagmins are calcium sensors localized in the β-cell insulin granules, and are required for vesicle fusion and glucose-stimulated insulin release (*38–40*). *SYT13* is a member of the synaptotagmin family and is predicted to be involved in calcium-regulated exocytosis. *SYT13* is down-regulated in human T2D islets and silencing of *SYT13* impairs insulin secretion in INS1-832/13 cells (*41*). To determine whether down-regulation of *CACNA1A* and *SYT13* seen in *HNF1A* mutant cells (**Fig. 1H**) reproduced the reduced intracellular calcium levels and reduced glucose-stimulated insulin secretion, we generated hESC *CACNA1A* KO and *SYT13* KO cell lines using CRISPR/Cas9. Sanger sequencing revealed homozygous mutations resulting in frame shifts (**Fig. S8C**). Differentiation of hESC *CACNA1A* KO and hESC *SYT13* KO generated INS-GFP organoids (**Fig. 3E**) comprised of ∼51% PDX1/CPEP and ∼16% GCG positive cells (**Fig. 3F and S8D**) with no differences compared to isogenic WT cells. Dispersed hESC *CACNA1A* KO *β*-like cells (**Fig. S8E**) had significantly reduced intracellular calcium levels compared to hESC WT cells (**Fig. 3G and S8F**). The reduction of intracellular calcium in hESC *CACNA1A* KO was intermediate to hESC *HNF1A* KO cells, indicating that *CACNA1A* is not the only gene affecting intracellular calcium levels in hESC *HNF1A* KO cells. Intracellular calcium levels in *SYT13* KO where unchanged compared to WT cells (**Fig. 3G and S8F**). Reduced basal and stimulated insulin secretion were observed in both hESC *CACNA1A* KO and hESC *SYT13* KO sc*β*-like cells at intermediate levels to *HNF1A* KO sc*β*-like cells (**Fig. 3H**). Treatment with tolbutamide or KCl stimulated insulin secretion in all KO lines comparable to the corresponding WT lines. Impairments in insulin release due to reduced levels of these molecules can be overcome by elevating calcium beyond physiological levels using tolbutamide or KCl. The milder phenotypes of cells deficient for single target genes compared to *HNF1A* KO cells suggests that the combined effects on several HNF1A target genes may be responsible for reduced glucose-stimulated insulin secretion.

### *HNF1A* deficiency alters the stoichiometry of insulin to C-peptide secretion

To interrogate the function of hESC *HNF1A* KO *β*-cells *in vivo*, we monitored circulating human insulin and C-peptide concentrations in euglycemic mice transplanted with sc-islet-like clusters. By 4 weeks post-transplantation, plasma human C-peptide concentrations were significantly lower in mice transplanted with hESC *HNF1A* KO *β*-cells compared to hESC *HNF1A* WT *β*-cells (**Fig. 4A**). Circulating human C-peptide concentrations in mice transplanted with hESC *HNF1A* WT β-cells increased over time, reaching human physiological levels (652±146 pM) 24-30 weeks post-transplantation. In mice transplanted with hESC *HNF1A* KO *β*-cells, plasma human C-peptide concentrations increased to a maximum of 352±142 pM at 30 weeks post-transplantation (**Fig. 4A**). In mice transplanted with hESC *HNF1A* WT *β*-cells, human insulin was detected as early as 4 weeks post-transplantation and concentrations increased over time, reaching human physiological levels (12.74±2.1 mU/L) 24-30 weeks post-transplantation. However, mice transplanted with hESC *HNF1A* KO *β*-cells had virtually undetectable plasma human insulin concentration for 30 weeks (**Fig. 4B**) despite high plasma human C-peptide levels (>300 pM). These differences in hormone secretion between *HNF1A* mutant and *HNF1A* WT scβ-cells were not due to reduced scβ-cell mass *in vivo* (**Fig. 2I**). These findings *in vivo* led us to quantify the secretion ratio of those hormones in sc*β*-cells *in vitro*. For equivalent secretion of human C-peptide at basal glucose condition (**Fig. S8G**), secretion of human insulin was significantly reduced in hESC *HNF1A* KO lines (**Fig. S8H**), leading to decreased insulin:C-peptide secretion ratios *in vitro* (**Fig. S8I**). Thus, the absence of insulin in the plasma was not a limitation of the sensitivity of the assay but a defect in the stoichiometry of insulin to C-peptide in circulation. While plasma of mice grafted with hESC *HNF1A* WT *β*-cells consistently showed an insulin:C-peptide molar ratio of 0.22±0.11 from week 4 to 30 post-transplantation, mice grafted with hESC *HNF1A* KO β-cells had an 18-fold lower ratio (**Fig. 4C**). This decrease in circulating insulin:C-peptide ratio was reciprocal to a 3-fold increase in the insulin:C-peptide ratio of intracellular content from hESC *HNF1A* KO isolated grafts (**Fig. 4D**), demonstrating complementary imbalance of insulin to C-peptide. The altered insulin:C-peptide ratio was not due to differences in insulin processing since insulin:proinsulin ratios in hESC *HNF1A* KO and hESC *HNF1A* WT grafts were identical *in vivo* (**Fig. 4E**) and *in vitro* (**Fig. S8J-S8L**) and no differences were found in the transcript levels of processing genes (*PC1*/*PC3*) in RNA sequencing analysis (**Table S3-S5**). Impaired stoichiometry of circulating insulin to C-peptide was also observed in mice transplanted with two additional homozygous mutant hESC lines (**Fig. S8M**). Therefore, *HNF1A* deficiency not only impairs insulin secretion, but also the stoichiometry of circulating insulin to C-peptide ratios.

**Fig. 4.**
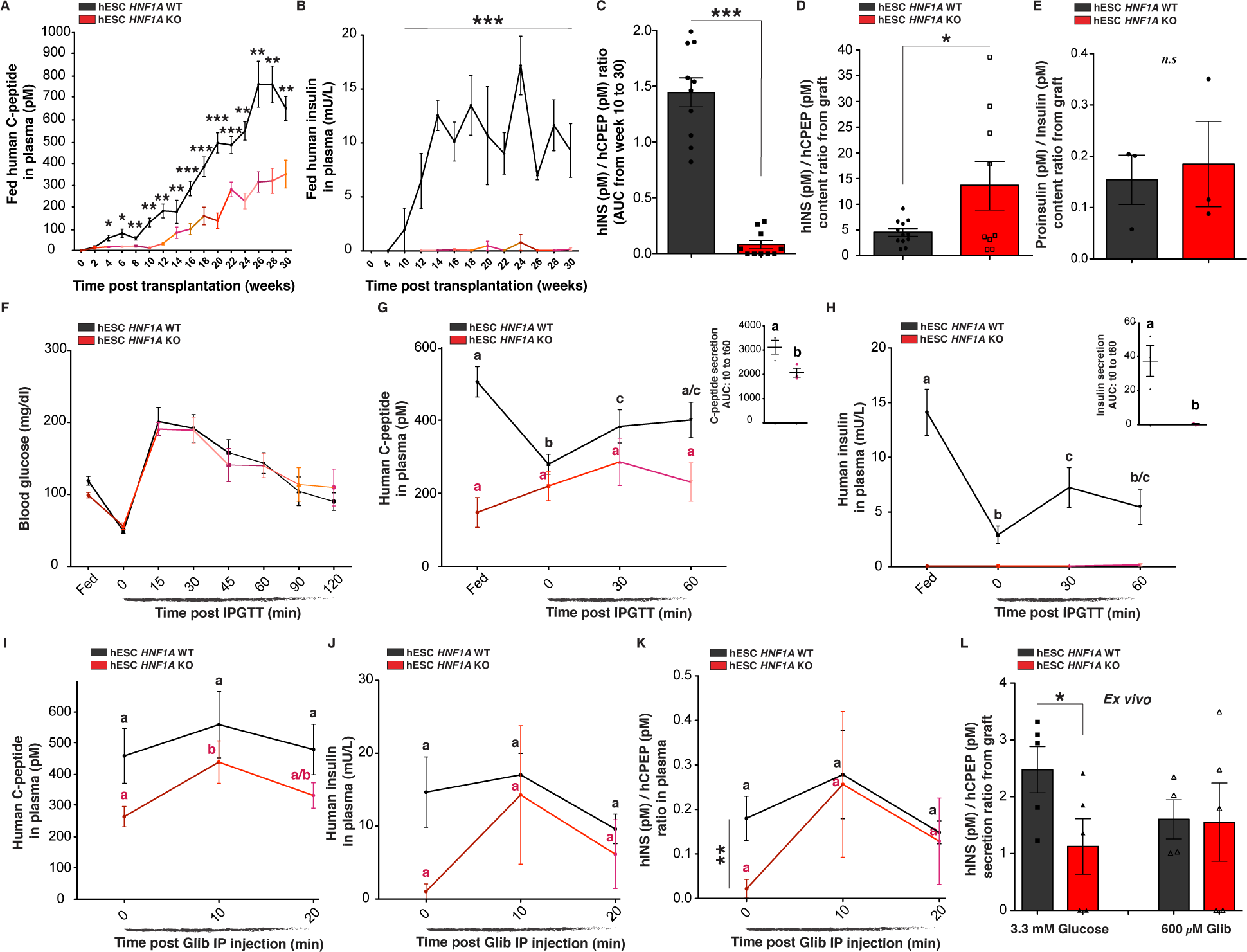
*HNF1A* deficiency alters the stoichiometry of insulin to C-peptide secretion *in vivo*. **(A)** Plasma human C-peptide (pM) and **(B)** human insulin secretion (mU/L) in plasma of *ad libitum-fed* mice transplanted with hESC-derived endocrine cells (*HNF1A* WT n=18 and KO n=7). **(C)** hINS(pM)/hCPEP(pM) ratios displayed as area under the curve (AUC) from week 10 to week 30 post-transplantation. (**D and E**) Content ratio from isolated grafts: **(D)** hINS(pM)/hCPEP(pM) content ratio and **(E)** Proinsulin(pM)/Insulin(pM) content ratio. (**F-H**) IPGTT in mice transplanted with hESC-derived endocrine cells (*HNF1A* WT n=10 and KO n=8) in *ad libitum-fed* state and during an iPGTT (t0, t30 and t60). **(F)** Blood glucose concentrations (mg/dl), **(G)** human C-peptide (pM) and **(H)** human insulin secretion (mU/L) in plasma. AUC: area under the curve from t0 to t60. p-values were b: p<0.001, c: p<0.05, a/c: p<0.05 and b/c: p<0.01; two-tailed t-test. (**I-K**) Glibenclamide injection in *ad libitum-fed* mice transplanted with hESC-derived endocrine cells (*HNF1A* WT n=6 and KO n=5). **(I)** Human C-peptide secretion (pM), **(J)** human insulin secretion (mU/L) and **(K)** hINS (pM)/hCPEP (pM) secretion ratios. p-values were b: p<0.05. **(L)** hINS(pM)/hCPEP(pM) secretion ratios *ex vivo* in response to indicated secretagogues. All mice were transplanted with clusters of hESC-derived endocrine cells at day 27 of differentiation, and grafts were isolated 30 weeks post-transplantation for *ex vivo* analysis. All protein concentrations were measured by ELISA. (n) represents the number of biological replicates. For scatter plots, each point in plots represents an independent biological experiment (n). Data are represented as mean ± SEM. Different letters designate significant differences within group. p-values: *p<0.05, **p<0.01, ***p<0.001; two-tailed t-test. n.s: non-significant. See also Fig. S8.

To evaluate glucose-stimulated insulin secretion, we performed an intraperitoneal glucose tolerance test (iPGTT) in transplanted mice. During fasting and following an iPGTT, mice transplanted with hESC *HNF1A* WT β-cells displayed normal human insulin and C-peptide secretion profiles (**Fig. 4F-4H**). In contrast, hESC *HNF1A* KO β-cells were non-responsive to glucose: plasma human C-peptide was not decreased by fasting and did not increase after glucose injection with concentrations significantly lower than in mice transplanted with hESC *HNF1A* WT β-cells (**Fig. 4G**). Remarkably, circulating human insulin remained undetectable in animals engrafted with hESC *HNF1A* KO β-cells despite high plasma levels of human C-peptide (**Fig. 4H**). Therefore, *HNF1A* deficiency affects glucose-stimulated insulin secretion, and C-peptide is released in a constitutive manner independent of metabolic state, resulting in an altered ratio of insulin to C-peptide.

### Glibenclamide restores insulin secretion in *HNF1A* deficient scβ-cells

In contrast to glucose, intra-peritoneal injection of the sulfonylurea drug, glibenclamide, in mice transplanted with hESC *HNF1A* KO β-cells showed a significant increase in C-peptide 10 minutes after glibenclamide injection (**Fig. 4I**). Surprisingly, human insulin concentrations in mice transplanted with hESC *HNF1A* KO β-cells increased from undetectable levels to 14±21 mU/L, reaching levels comparable to those in the control group (**Fig. 4J**). Insulin:C-peptide ratios in plasma of animals transplanted with hESC *HNF1A* KO β-cells were increased 10-fold by glibenclamide, equal to the ratios in mice transplanted with hESC *HNF1A* WT β-cells (**Fig. 4K**). Clearance of insulin from the circulation occurred with the same kinetics in mice grafted with hESC *HNF1A* WT and hESC *HNF1A* KO β-cells 20 minutes after glibenclamide administration (**Fig. 4K**), excluding insulin stability or clearance in plasma as the cause for the low insulin levels in mice transplanted with *HNF1A* mutant scβ-cells. To further test whether the reduced insulin:C-peptide secretion ratios from hESC *HNF1A* KO β-cells were due to secretory defects and not insulin stability or clearance, we isolated the hESC *HNF1A* KO grafts from mice and identified a significant decrease in insulin:C-peptide secretion ratios compared to *HNF1A* WT grafts *ex vivo*. This ratio was restored after exposure of the grafts to glibenclamide (**Fig. 4L**).

Thus, membrane depolarization initiated by closure of the ATP-sensitive K^+^ (K_ATP_) channels (target of glibenclamide) and high intracellular calcium levels allow the secretion of insulin granules that are abnormally retained in *HNF1A* mutant *β*-cells.

### *HNF1A* deficiency causes abnormal insulin granule structure

To identify the basis of the insulin secretory defect in hESC *HNF1A* KO β-cells we examined insulin granule morphology by electron microscopy from 30-week explants of normoglycemic mice. hESC *HNF1A* WT grafts were comprised of scβ-cells with a characteristic high-electron density core separated from limiting membranes by a “halo” (**Fig. 5A and S9A**). These morphologies are consistent with electron microscopy of normal human β-cells. In contrast, hESC *HNF1A* KO islets-like were comprised of scβ-cells where the majority (>90%) of the insulin granules were abnormally enlarged (**Fig. 5A-5C**) with a diffuse electron-light insulin core (**Fig. 5D-5E and S9A**), characteristic of immature secretory granules. Enlarged insulin granules were further confirmed by immunogold staining for human C-peptide (**Fig. 5F**). Similar results were observed *in vitro*, where hESC *HNF1A* mutated cells had increased insulin granule diameter with a diffuse light insulin core (**Fig. S9B and S9C**). For sc*α*-cells, no differences in morphology, structure or granule size between *HNF1A* genotypes was seen (**Fig. S9D**). While these abnormal insulin granules are not secreted in response to glucose, they can be secreted in response to sulfonylureas. These results show that *HNF1A* is required for normal insulin granule formation and function.

**Fig. 5.**
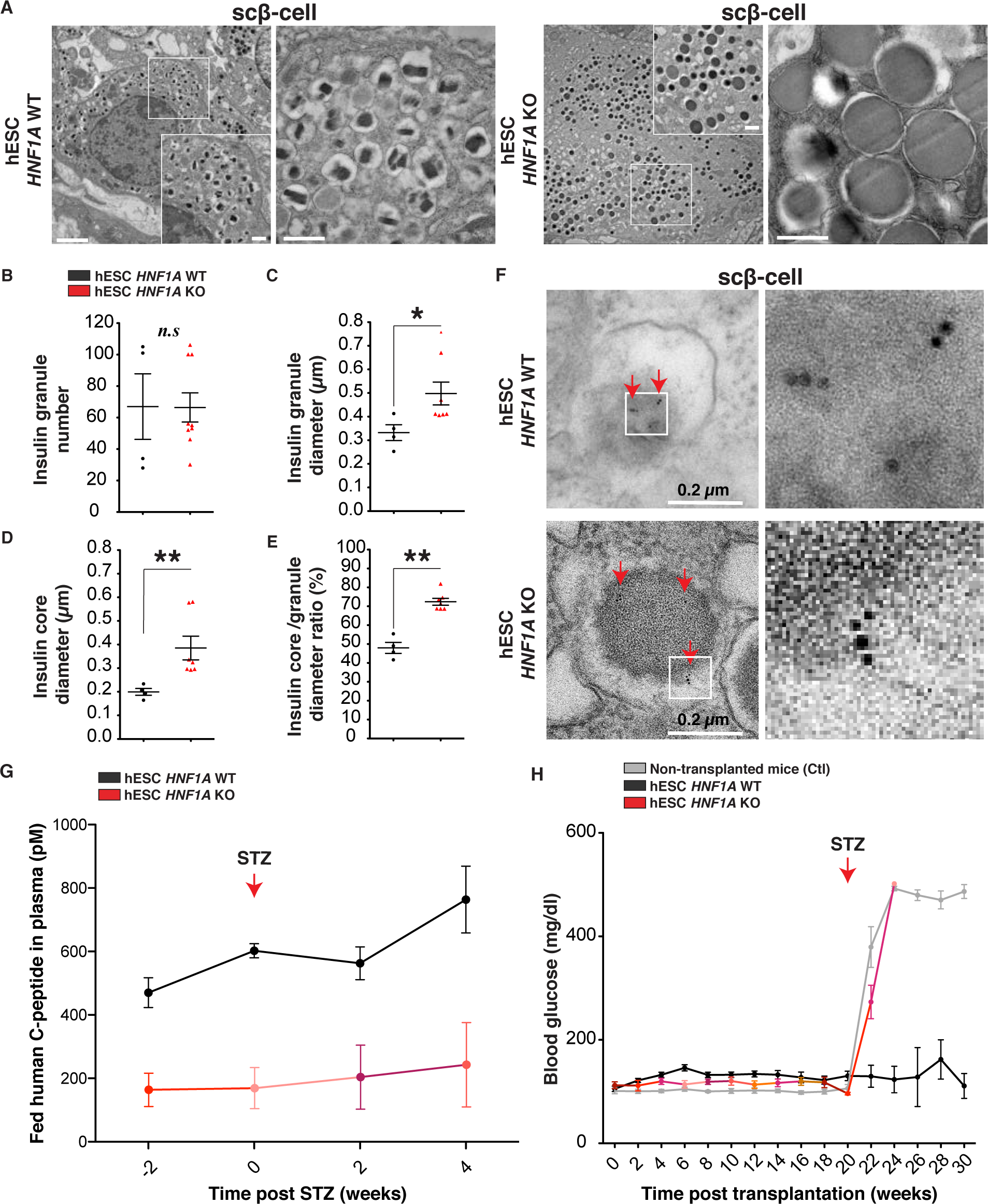
*HNF1A* deficiency causes abnormal insulin granule structure and fails to maintain glucose homeostasis in diabetic mice. (**A and B**) Electron microcopy (EMC) analysis of isolated grafts 30 weeks post-transplantation for hESC-derived endocrine cells (*HNF1A* WT and KO). Explants are from euglycemic mice. **(A)** Representative EMC images of scβ-cells with (**B**) quantification of insulin granule number, **(C)** insulin granule diameter (µm), **(D)** insulin granule core diameter (µm) and **(E)** insulin granule core diameter to insulin granule diameter ratio (%) per cell. Each point in plots is the average of insulin granules per scβ-like-cells. **(F)** Representative EMC images of scβ-cells with human C-peptide immunogold labelling. Scale bars: 2 µm in low and 0.5 µm in high magnification. **(G)** Fed human C-peptide (pM) in plasma of *ad libitum-fed* mice transplanted with hESC-derived endocrine cells (*HNF1A* WT n=8 and KO n=3) measured weeks before (-2) and after STZ treatment (0, 2 and 4). **(H)** Blood glucose concentrations (mg/dl) in *ad libitum-fed* mice transplanted with hESC-derived endocrine cells (*HNF1A* WT n=18 and KO n=10) or without cells (Ctl n=14). Mice (*HNF1A* WT n=6, KO n=3 and Ctl n=5) were injected (red arrow) with streptozotocin (STZ). All mice were transplanted with hESC-derived endocrine cells at day 27 of differentiation, and grafts were isolated 30 weeks post-transplantation for *ex vivo* analysis. All protein concentrations were measured by ELISA. (n) represents the number of biological replicates. For scatter plots, each point in plots represents an independent biological experiment (n). Data are represented as mean ± SEM. p-values: *p<0.05, **p<0.01, ***p<0.001; Mann-Whitney test. n.s: non-significant. See also Fig. S9 and S10.

### *HNF1A* deficient scβ-cells are unable to maintain glucose homeostasis in diabetic mice

To test the ability of hESC *HNF1A* KO β-cells to maintain *in vivo* normoglycemia, endogenous murine β-cells were ablated by the administration of streptozotocin (STZ) 22 weeks post-transplantation. STZ is a drug that, by virtue of species-related difference in glucose transporters, specifically targets murine β-cells without affecting human β-cells (*42*). We first confirmed that *HNF1A* deficiency did not increase sensitivity of human cells to STZ. There was no change in cell death (**Fig. S10A**) or C-peptide secretion (**Fig. S10B**) when *HNF1A* WT or KO scβ-like-cells were exposed to STZ *in vitro*. *In vivo*, circulating plasma human C-peptide from mice transplanted with *HNF1A* WT or *HNF1A* KO cells was unchanged or increased post STZ treatment (**Fig. 5G**). Mouse C-peptide was undetectable by ELISA (**Fig. S10C**) and immunohistochemistry of the pancreas (**Fig. S10D**) demonstrating successful and specific ablation of mouse β-cells. Mice transplanted with the hESC *HNF1A* WT β-cells were normoglycemic (∼100 mg/dl and HbA1C ∼5%) for at least 10 weeks post STZ injection (**Fig. 5H and S10E**) and had normal human insulin and C-peptide secretion profiles during an iPGTT (**Fig. S10F-S10H**). In contrast, mice transplanted with hESC *HNF1A* KO β-cells did not increase human C-peptide secretion (**Fig. 5G**) after becoming severely diabetic (blood glucose >500 mg/dl) within one week following STZ injection (**Fig. 5H**). Consistent with *in vitro* studies, hESC *HNF1A* KO β-cells are unresponsive to blood glucose even at high levels, failing to maintain systemic glucose homeostasis in a mouse model of β-cell deficient diabetes.

### *HNF1A* R200Q homozygous mutation is pathogenic and causes a developmental bias towards the *α*-cell fate *in vitro*

These studies demonstrate the molecular, cellular and functional consequences of complete *HNF1*A deficiency, a situation that does not occur clinically, but was created experimentally to amplify the effects of HNFIA functional hypomorphism in order to enable mechanistic studies. MODY3 patients are heterozygous for the causal *HNF1A* alleles; to more directly characterize the consequences of heterozygous patient-specific mutations in *HNF1A*, we differentiated two heterozygous MODY3 iPSC lines (+/460_461insCGGCATCCAGCACCTGC and +/R200Q), isogenic mutation-corrected MODY3 iPSC lines (R200Q corrected: WT) and hESC *HNF1A* Het line into endocrine cells. Heterozygous mutations in *HNF1A* did not affect the generation of definitive endoderm cells (SOX17^+^) at day 3 of differentiation or pancreatic progenitor cells (PDX1^+^/NKX6.1^+^) at day 11 of differentiation compared to WT lines as determined by immunohistochemistry (**Fig. S11A**). At the endocrine stage (day 27), cell clusters or organoids were morphologically indistinguishable and all heterozygous cell lines differentiated efficiently to all pancreatic endocrine cells (sc*α*-, sc*β*- and scδ-like cells) and sc*β*-like cells co-expressed PDX1/NKX6.1/HNF1A (**Fig. S11B**). HNF1A protein was detected in WT, MODY3 R200Q Het (+/R200Q) and MODY3 R200Q-corrected WT (+/+) iPSC-derived endocrine cells by western blot (**Fig. S11C**). In contrast to *HNF1A* truncated lines, the heterozygous point mutation R200Q of MODY3 iPSC line did not affect the total amount of HNF1A protein, suggesting that the mutant R200Q protein was produced (**Fig. S11C**). No significant differences were found in endocrine cell types (sc*α*-, sc*β*- and scδ-like cells) between *HNF1A* WT and heterozygous cell lines at day 27 of differentiation by immunohistochemistry (**Fig. S11D**).

To determine the consequences of heterozygosity for *HNF1A* in our MODY3 patient lines for the bias towards *α*-cell fate, we measured the percentage of glucagon cells co-expressing C-peptide in two MODY3 iPSC Het (+/460ins and +/R200Q) lines *in vitro*. We found no significant differences between MODY3 iPSC lines and control iPSC R200Q-corrected WT lines (**Fig. S11E**). To determine the pathogenicity of the *HNF1A* R200Q mutation, we knocked out of the *HNF1A* wildtype allele in a MODY3 iPSC Het line (R200Q/-), which resulted in a significant ∼70% increase in GCG and CPEP co-expressing cells compared to the R200Q-corrected WT control lines (**Fig. S11E and S11F**). We also detected a significant increase in the percentage of GCG and CPEP co-expressing cells in hESC *HNF1A* R200Q homozygous (R200Q/R200Q) and hESC *HNF1A* Het (+/-) lines compared to hESC WT (+/+) lines (**Fig. S11G and S11H**). Thus, while both heterozygous MODY3 patient cells (+/460ins and +/R200Q) show no developmental bias towards the *α*-cell fate, complete loss of *HNF1A* WT allele in MODY3 iPSC R200Q (R200Q/-) and hESC *HNF1A* R200Q homozygous (R200Q/R200Q) result in a stronger developmental bias towards *α*-cell fate, indicating the pathogenicity of R200Q mutation. Interestingly, the hESC heterozygous line harboring a frameshift mutation in the DNA-binding domain and causing a premature stop codon presented a stronger bias towards *α*-cell fate. Previous studies in MODY3 patients have shown that frameshift mutations in the DNA-binding domain cause dominant-negative effects affecting the DNA binding capacity of the normal HNF1A (*43*).

### *HNF1A* haploinsufficiency gradually impairs scβ-cell functional capacity in the context of metabolic stress

In order to assess the developmental potential of MODY3 patient-specific mutations *in vivo*, we transplanted pancreatic islet-like clusters derived from iPSCs using identical methods as previously described. While 87.9% (51/58) of mice transplanted with isogenic MODY3 iPSC Het lines (+/R200Q) and R200Q-corrected WT (+/+) were teratoma-free; the majority (61.5%, 8/13) of the MODY3 iPSC Het (460ins) and control iPSC WT lines showed teratoma formation (**Fig. S11I**). Because of variable teratoma formation among different iPSC lines (*20*), only teratoma-free mice were used for further analysis. Teratoma-free transplanted mice had 67.3% (35/52) engraftment efficiency. Thirty weeks post-transplantation, we isolated grafts from normoglycemic animals (**Fig. S11J**) and found no significant differences in glucagon-to-insulin content ratios or endocrine cell types between MODY3 iPSC Het (+/R200Q) and MODY3 iPSC R200Q-corrected WT (+/+) control grafts (**Fig. S11K and S11L**). These results are consistent with our iPSC *in vitro* results; however, a recent study from cadaveric MODY3 human islets with a T260M heterozygous mutation in DNA-binding domain (*29*), showed a bias of endocrine cells toward the *α*-cell fate (**Fig. 2K**). The same phenotype was observed in hESC *HNF1A* heterozygous lines with frameshift mutation in the DNA-binding domain **(Fig. S11H)**. Mutations affecting the DNA-binding domain or truncating mutations are associated with earlier onset of diabetes in MODY3 patients (*44*). Thus, different mutations affect HNF1A function to different degrees, resulting in different phenotypes in patients as well as in cellular phenotypes in a stem cell system.

To understand the function of MODY3 iPSC Het *β*-cells in a physiological context, we monitored circulating human insulin and C-peptide in transplanted euglycemic mice over 30 weeks. Mice transplanted with iPSC-derived cells from heterozygous MODY3 patient (+/460_461insCGGCATCCAGCACCTGC) had low, but detectable circulating human C-peptide two, four, eight (**Fig. S12A**) and thirty weeks post transplantation (**Fig. S12B**) with undetectable human insulin (**Fig. S12C**). Since human insulin levels are under the minimal threshold limit for detection by ELISA, this deficiency can be attributable to the lower ability of these mutant iPSCs to generate functional insulin producing cells *in vivo*. In mice transplanted with iPSC-derived cells from MODY3 patient (+/R200Q) and R200Q-corrected WT (+/+) isogenic lines, circulating human C-peptide (**Fig. S12B**) and human insulin (**Fig. S12C**) concentrations reached similar levels 30 weeks post-transplantation. Transplantation of mice with hESC *HNF1A* Het (+/-) and hESC *HNF1A* R200Q homozygous (R200Q/R200Q) line resulted in low circulating concentrations of human C-peptide over thirty weeks post transplantation (**Fig. S12A and S12B**) with low to undetectable human insulin in hESC *HNF1A* Het (+/-) and hESC *HNF1A* Hom (R200Q/R200Q) as compared to WT lines (**Fig. S12C**), thus affecting insulin:C-peptide secretion ratios. These results show that *HNF1A* heterozygous frameshift mutations in the DNA-binding domain (hESC +/-) and *HNF1A* R200Q homozygosity (hESC R200Q/R200Q) are pathogenic and that in the heterozygous point mutations (iPSC +/R200Q), it is less detrimental to *β*-cell function.

To further interrogate the consequence of the R200Q mutation in *β*-cells function and evaluate glucose-stimulated insulin secretion, we performed an intraperitoneal glucose tolerance test (iPGTT) in mice transplanted with the MODY3 isogenic lines (R200Q Het and R200Q-corrected WT), as these iPSC lines allowed the reliable generation of transplanted mice with high C-peptide and insulin levels over thirty weeks post transplantation (**Fig. S12D and S12E**). During an iPGTT, mice transplanted with both isogenic lines showed comparable changes in blood glucose (**Fig. S12F**) and plasma human C-peptide (**Fig. 6A**); plasma human insulin concentrations dropped below detection during fasting and increased upon glucose injection (**Fig. 6B**), demonstrating homeostatic glucose responsiveness. Glibenclamide injection increased C-peptide and insulin concentrations in both genotypes (**Fig. S12G and S12H**) at equivalent ratios (**Fig. S12I**). No significant differences in insulin secretion (**Fig. S12J**) or endocrine hormone content were found *in vitro* or *in vivo* between genotypes (**Fig. S12K and S12L**).

**Fig. 6.**
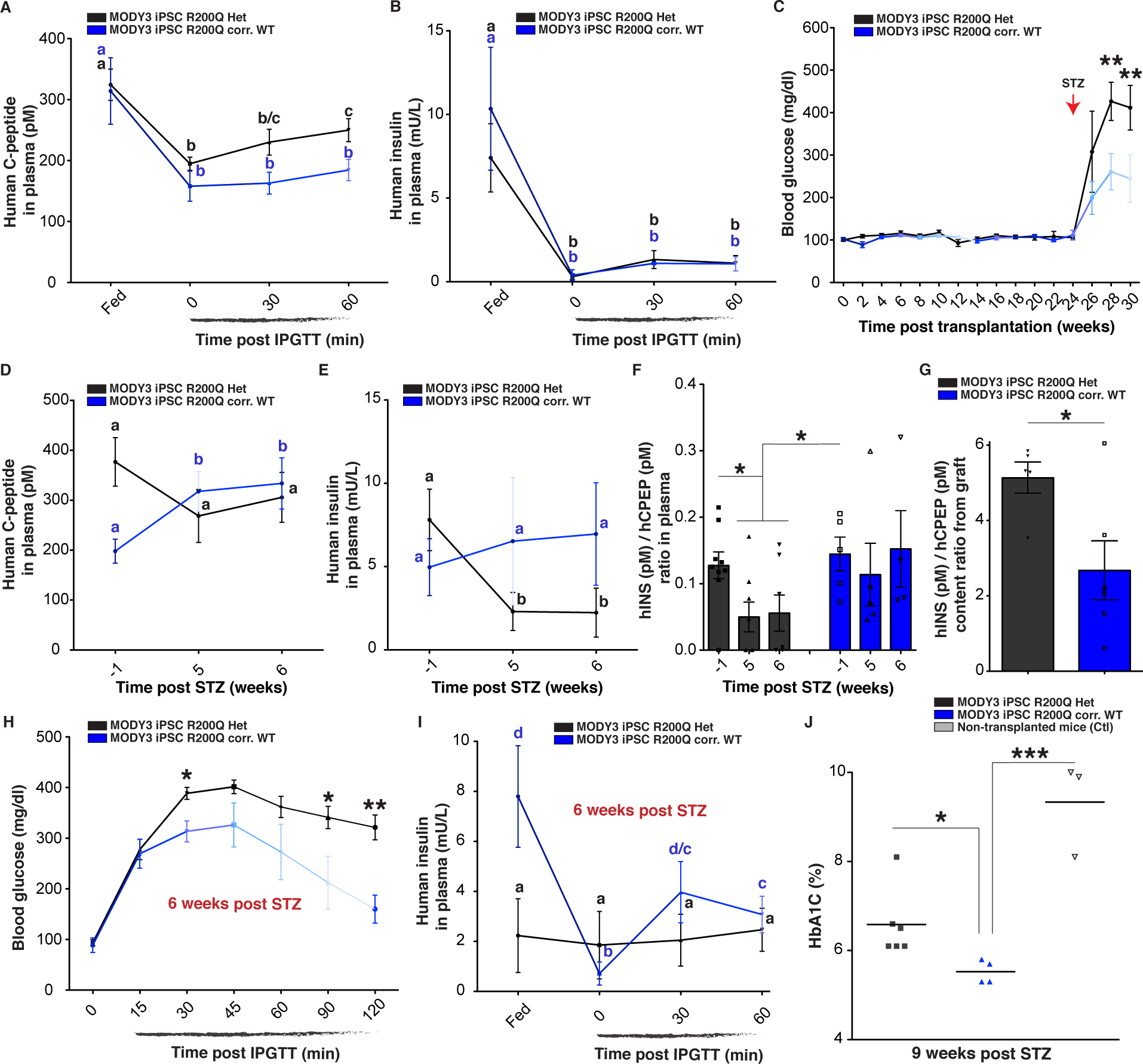
*HNF1A* haploinsufficiency gradually impairs scβ-cell function *in vivo*. (**A-C**) Human C-peptide (pM) and human insulin levels in plasma of *ad libitum-fed* mice transplanted with MODY3 iPSC-derived endocrine cells (*HNF1A* +/R200Q and R200Q-corrected WT). **(A)** Plasma human C-peptide (pM) and **(B)** human insulin secretion (mU/L) in mice transplanted with MODY3 iPSC-derived endocrine cells (R200Q Het n=10 and R200Q corr. WT n=5) in *ad libitum-fed* state and during an iPGTT (t0, t30 and t60). p-values were b: p<0.001, b/c: p<0.01 and c: p<0.05. **(C)** Blood glucose concentrations (mg/dl) in *ad libitum-fed* mice transplanted with MODY3 iPSC**Human** derived endocrine cells (R200Q Het n=21 and R200Q corr. WT n=32). Mice (R200Q Het n=8, R200Q corr. WT n=8 and Ctl n=5) were injected (red arrow) with STZ. **(D)** Human C-peptide (pM), **(E)** human insulin (mU/L) (R200Q Het n=9 and R200Q corr. WT n=6), and **(F)** hINS (pM)/hCPEP (pM) secretion ratios monitored in plasma of *ad libitum-fed* mice weeks before (-1) and after (5 and 6) STZ treatment. p-values were b and d: p<0.01; c: p<0.05. **(G)** Quantification of hINS(pM)/hCPEP(pM) content ratios from isolated grafts. (**H-I**) IPGTT in mice transplanted with MODY3 iPSC-derived endocrine cells (R200Q Het n=5 and R200Q corr. WT n=4) 6 weeks post STZ treatment in *ad libitum-fed* state and during an iPGTT (t0, t30 and t60). **(H)** Blood glucose concentrations (mg/dl) and **(I)** human insulin secretion (mU/L) in plasma. p-values were b p<0.01, d/c and c: p<0.05. (R200Q Het n=5 and R200Q corr. WT n=4). **(J)** HbA1C (%) in mice transplanted with MODY3 -iPSC-derived endocrine cells or non-transplanted (Ctl) 9 weeks after STZ treatment. All mice were transplanted with iPSC-derived endocrine cells at day 27 of differentiation, and grafts were isolated 30-35 weeks post-transplantation for *ex vivo* analysis. All protein concentrations were measured by ELISA. For scatter plots, each point in plots represents an independent biological experiment (n). Data are represented as mean ± SEM. Different letters designate significant differences within groups. p-values were *p<0.05, **p<0.01, ***p<0.001; two-tailed t-test. n.s: non-significant. See also Fig. S11-S13.

MODY3 patients generally display normal glucose tolerance during early childhood and exhibit symptomatic diabetes only in their late teens or early adulthood (*9*) depending on the type and position of the *HNF1A* mutation (*44*), indicating that progression of the disease occurs over several years and potentially involves metabolic stressors. To determine whether there are functional differences between heterozygous MODY3 mutated (+/R200Q) and MODY3 R200Q-corrected WT (+/+) scβ-cells, we treated mice with STZ, thereby exposing transplanted scβ-cells to higher insulin demand due to ablation of mouse endogenous β-cells. Two weeks post STZ, animals transplanted with MODY3 iPSC R200Q Het (+/R200Q) cells showed rapid progression to hyperglycemia (**Fig. 6C**). The increase in blood glucose from those mice (n=9) was accompanied by a failure to increase circulating human C-peptide (**Fig. 6D**) and a significant reduction of human insulin levels (**Fig. 6E**) over 5-6 weeks after STZ treatment. In contrast, in mice (n=6) transplanted with isogenic MODY3 iPSC R200Q-corrected WT cells (+/+), concentrations of human plasma C-peptide (**Fig. 6D**) and insulin increased (**Fig. 6E**).

Five to six weeks post STZ treatment, we found that the ratio of circulating insulin to C-peptide fell by 55% in mice transplanted with MODY3 iPSC R200Q Het (+/R200Q) cells. In contrast, insulin:C-peptide ratios remained constant in mice transplanted with MODY3 R200Q-corrected WT (+/+) cells (**Fig. 6F**). The decrease in circulating insulin:C-peptide ratios was reciprocal to a significant increase by 74% in the insulin:C-peptide ratios of intracellular content from isolated *HNF1A* mutated grafts compared to isogenic controls (**Fig. 6G**), demonstrating complementary imbalance of insulin to C-peptide secreted. These results show that *HNF1A* R200Q haploinsufficiency gradually impairs the stoichiometry of circulating insulin:C-peptide, and that gene correction of the R200Q mutation protects MODY3 iPSC β-cells from acquiring this imbalance.

We then performed iPGTTs two, four, six and eight weeks post STZ administration. MODY3 iPSC R200Q Het (+/R200Q) scβ-cells showed progressive impairment of glucose-stimulated insulin secretion: mild at two weeks post STZ (**Fig. S13A and S13B**) and completely unresponsive insulin release to glucose challenges four weeks post STZ (**Fig. S13C and S13D**). Mice transplanted with MODY3 scβ-cells fail to clear glucose from the circulation during a glucose tolerance test at six to eight weeks post STZ treatment (**Fig. 6H and S13E**). *HNF1A* R200Q heterozygous (+/R200Q) sc*β*-cells had no glucose-stimulated insulin secretion, whereas MODY3 R200Q-corrected WT (+/+) scβ-cells were sensitive to fasting, remained glucose responsive six to eight weeks post STZ treatment, and cleared glucose from circulation (**Fig. 6H-6I and S13E-S13F**). Mice transplanted with MODY3 iPSC R200Q Het (+/R200Q) scβ-cells became diabetic as shown by elevated blood glucose (**Fig. S13G**) and HbA1c levels (**Fig. 6J**) compared to control mice. These results show that MODY3 iPSC β-cells (+/R200Q) fail to compensate for higher insulin demands, and that the hyperglycemia is due to gradual development of insulin secretory defects, characterized by a disruption of the stoichiometry of insulin and constitutive C-peptide release. Correction of R200Q mutations protects MODY3 iPSC β-cells from acquiring these abnormal insulin secretion profiles.

### MODY3 patients have low non-fasting insulin to C-peptide ratios in plasma

To determine whether discordant concentrations of circulating insulin and C-peptide also occurred in MODY3 patients, we measured islet endocrine hormones in the plasma of nine MODY3 patients (**Table S6**). MODY3 patients were divided into three therapeutic groups: no sulfonylurea treatment (5/9 patients); receiving sulfonylurea treatment (3/9 patients), and receiving Sitagliptin (GLP1 agonist) treatment (1/9 patients). In the group without sulfonylurea treatment, both MODY3 and MODY2 (hypomorphic mutations in glucokinase gene) patients had reduced non-fasting C-peptide levels in plasma compared to non-diabetic controls, 251±133 pM and 288±12 pM respectively (**Fig. 7A**). However, only MODY3 patients had undetectable human insulin levels (0 pM) in circulation, as opposed to MODY2 (14±8.2 pM) or non-diabetic controls (35±24 pM) (**Fig. 7B**). As in the stem cell model, this imbalance is apparent in an abnormal insulin:C-peptide secretion ratios in MODY3 patients (<0.002) as opposed to normal ratios in MODY2 patients (0.05±0.03) and non-diabetic controls (0.04±0.03) (**Fig. 7C**). In the group with sulfonylurea treatment, three MODY3 patients received glipizide. Those patients had non-fasting C-peptide concentrations of 616±301 pM (**Fig. 7A**) with circulating insulin of 25±12 pM (**Fig. 7B**) and normal insulin:C-peptide ratios (**Fig. 7C**), comparable to non-diabetic subjects. One MODY3 patient receiving Sitagliptin (dipeptidyl peptidase-4 inhibitor) also displayed an insulin:C-peptide ratio in plasma comparable to that of the controls. Patients who are insensitive to the insulinotropic effects of sulfonylureas, will respond clinically to GLP-1 agonists (*45*). There are multiple mechanisms by which GLP1R agonists may affect β-cell performance (*46*), and the findings in this subject are consistent with an effect of *HNF1A* deficiency on insulin secretory mechanisms that are glucose-dependent. No differences were found in the concentrations of non-fasting plasma glucagon in MODY3 patients compared to non-diabetic subjects (**Fig. 7D**). These results indicate that *HNF1A*, but not *GCK* mutations, affect the stoichiometry of circulating concentrations of insulin and C-peptide. The clinical efficacy of sulfonylureas in releasing insulin from β-cell granules and restoring normal insulin:C-peptide stoichiometry is consistent with our *in vitro* and *in vivo* studies of stem cell-derived islets, and with the clinical efficacy of sulfonylurea-type drugs in the treatment of MODY3 patients. The clinical differences of circulating hormone concentrations in MODY3 patients are not necessarily due entirely to effects on secretion rates, and could reflect genotype-related differences in the hepatic and renal handling of these peptides. However, genotype-specific stem cell-derived islets were grafted into genetically identical *HNF1A* (+/+) mice, and in those animals the alterations in the stoichiometry of circulating insulin and C-peptide are presumably determined solely by cell-autonomous characteristics of the grafted cells.

**Fig. 7.**
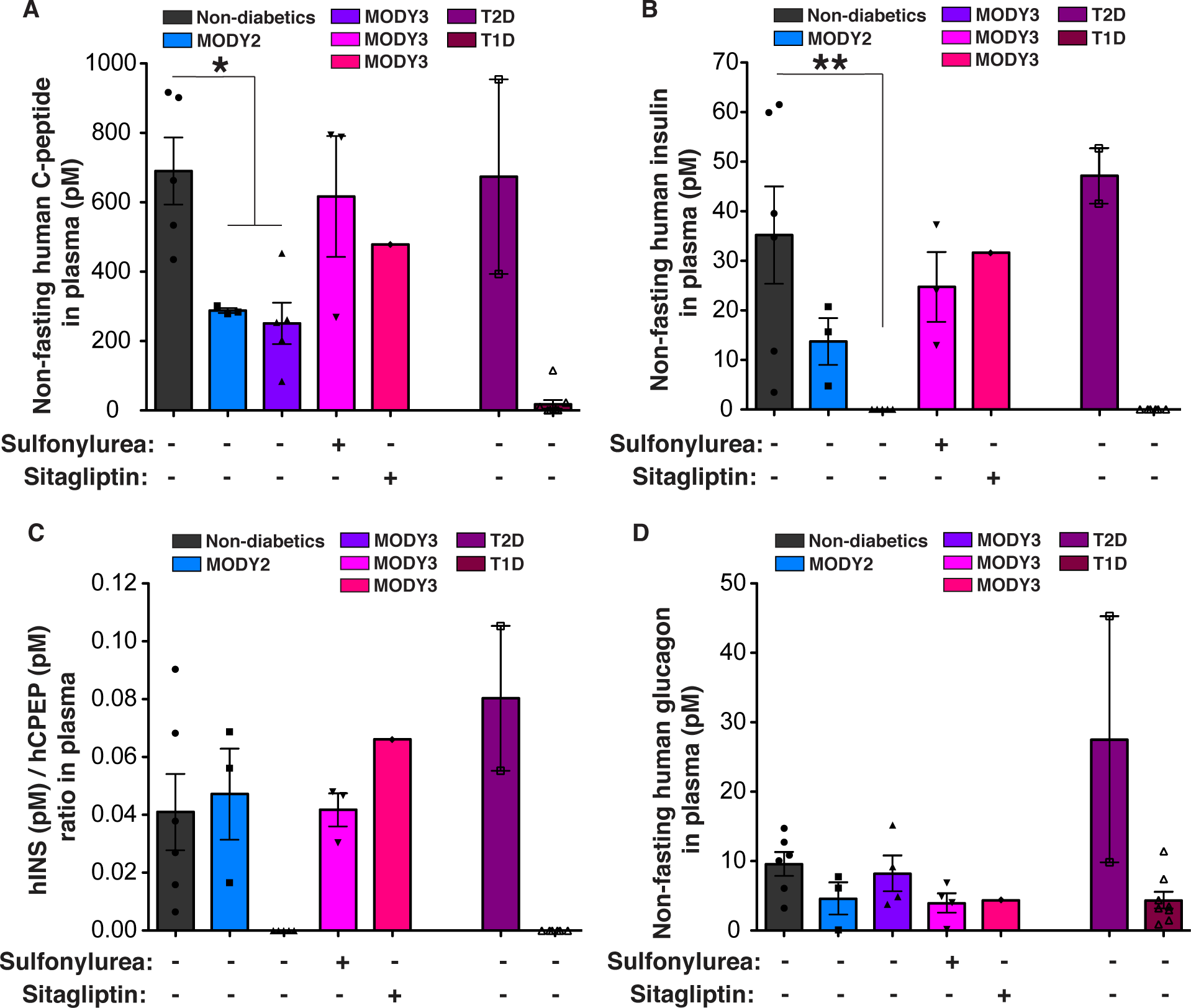
MODY3 patients have low non-fasting insulin to C-peptide ratios in plasma. (**A**) Clinical measurements of non-fasting human C-peptide (pM), (**B**) insulin (pM), (**C**) INS(pM)/CPEP(pM) ratio and (**D**) glucagon (pM) from plasma of non-diabetic subjects, MODY2 subjects, MODY3 subjects with (+) or without (-) drug treatment as indicated. Drugs taken chronically for treatment of impaired glucose metabolism. Additional controls are T2D subjects and T1D subjects. For scatter plots, each point in plots represents an independent biological experiment (n). Data are represented as mean ± SEM. p-values: *p<0.05, **p<0.01, ***p<0.001; Mann-Whitney test. See also Table S6.

## Discussion

Here we report the use of stem cell-derived islets containing *HNF1A* mutations to elucidate the molecular basis for apparent β-cell dysregulation of insulin secretion in *HNF1A* hESC null and MODY3 hypomorphic patient lines. Based on *in vitro* and *in vivo* developmental, morphological and functional assessments of stem cell-derived islets, as well as measurements of plasma concentrations of insulin and C-peptide in nine MODY3 patients, we conclude that *HNF1A* regulates endocrine cell fate, normal β-cell granule formation and function, including calcium-mediated exocytosis of insulin-containing granules, and the resultant stoichiometric secretion of insulin and C-peptide.

Hypomorphic mutations in *HNF1A* perturb expression of genes important in β-cell identity such as *PAX4*, leading to increases in glucagon gene expression, *α*-cell number and in the *α*:*β*-cell ratio in mature islets. This phenotype was observed in homozygous hESC *HNF1A* knockouts, but was not seen in MODY3 patient iPSC-derived heterozygous cells (+/460ins and +/R200Q). This increase was, however, reproduced in MODY3 patient iPSC- and hESC-derived cells rendered homozygous for R200Q (iPSC R200Q/- and in hESC R200Q/R200Q) indicating the pathogenicity of the R200Q allele. The hESC *HNF1A* line heterozygous for a frameshift mutation in the DNA-binding domain also showed the *α*-cell bias phenotype. Previous clinical studies in MODY3 patients have shown that the type and the position of *HNF1A* mutation is important in the timing of onset of the diabetic phenotype (*44*). The differences in the severity of the phenotype is likely due to a dominant-negative effect of frameshift mutations in the DNA-binding domain. Though DNA binding was not examined for the mutations studied here, previous studies have shown that frameshift mutations in the DNA-binding domain (c.422_423InsT) cause dominant-negative effects affecting the DNA binding capacity of the normal HNF1A molecule (*43*), as the mutant HNF1A will bind to the normal HNF1A via the intact dimerization domain in the N-terminus. A recent case study of cadaveric human islets from a 33-year old MODY3 patient (+/T260M mutation in the DNA-binding domain) with a 17-years history of diabetes also showed an increase in the ratio of *α*-cell to *β*-cells (*29*).

In *HNF1A* knockout stem cell-derived *β*-cells, glucose-stimulated insulin secretion is compromised in association with a reduction in *CACNA1A* and intracellular calcium levels, and down-regulation of *SYT13*, a gene involved in calcium-regulated granule exocytosis. We show that the administration of sulfonylurea increases intracellular calcium above physiological levels, enabling insulin vesicle fusion and secretion from *HNF1A* mutant *β*-cells. Consistent with this inference, the administration of sulfonylureas to MODY3 patients and mice transplanted with *HNF1A* knockout *β*-cells restores normal insulin secretion. *HNF1A* knockout cells accumulate enlarged insulin granules with diffuse electron-light insulin cores, which can be recruited by membrane depolarization in response to increases in intracellular calcium concentrations. Sulfonylureas may be stimulating release primarily from the more normal insulin granules and/or driving stoichiometric release of insulin and C-peptide from the enlarged, abnormal insulin granules. These results may be relevant to more prevalent forms of diabetes, including type 2 diabetes, in which reduced proportions of mature insulin granules have also been reported (*47, 48*).

Among some apparent *HNF1A* target genes, knockout of *PAX4*, *CACNA1A* and *SYT13* partially reproduced phenotypes seen in *HNF1A* deficient cells. As HNF1A orchestrates a network of genes involved at different aspects of β-cell development and function (*49*), a contribution of additional target genes to these phenotypes is likely. A recent report by Cardenas-Diaz et al. (*31*) identified *LINC01139* as an *HNF1A* target implicated in β-cell dedifferentiation and mitochondrial function. However, in *α*- or *β*-cells, we did not observe significant downregulation of *LINC01139* in *HNF1A* knockout or heterozygous cells.

The accumulation of abnormal insulin secretory granules in *HNF1A* knockout *β*-cells was associated with a loss of the 1:1 stoichiometric release of C-peptide and insulin from β-cells, characterized by constitutive secretion of C-peptide, and intracellular retention of insulin. Their equimolar secretion was restored by exposure of the cells to depolarization by sulfonylurea. Previous studies have identified a “constitutive-like” secretory pathway that, in immature β-cells, is characterized by accumulation of immature secretory granules that secrete newly synthesized C-peptide in molar excess of insulin (*50, 51*). The accumulation of abnormal insulin granules in *HNF1A* deficient β-cells is apparently associated with an increase in insulin granule membrane remodeling, as a result of which C-peptide is diverted most likely to the constitutive-like secretion pathway, accounting for reduced insulin to C-peptide ratios detected in the plasma of MODY3 patients and mice transplanted with *HNF1A* hypomorphic islets. Our data support this model, but more studies are necessary to validate this pathway and fully elucidate the contribution of secretory defects in MODY3 patients.

In cells with total *HNF1A* deficiency or knockout, uncoupling of C-peptide from insulin secretion is innate, while it is gradually acquired during progressive *β*-cell failure in heterozygous MODY3 patient cells. Measurement of plasma C-peptide has been widely and productively used in the study and clinical management of patients with diabetes (*52*). The observation in our study that C-peptide secretion can be dissociated from insulin secretion is clinically relevant when evaluating islet function in patients with MODY3. The ratio of insulin to C-peptide may actually be useful as a biomarker for progression of MODY3 and possibly other types of diabetes.

The insulin secretory phenotype of heterozygous MODY3 iPSC-derived *β*-cells was intermediate to that of cells with complete *HNF1A* deficiency. MODY3 iPSC-derived heterozygous *β*-cells (+/R200Q) initially functioned normally in transplanted mice, consistent with normal glucose tolerance of heterozygous carriers during early childhood, followed by symptomatic diabetes within the first three decades of life (**table S6** for age of onset) (*9*). Upon increasing the requirement for human insulin following ablation of the mouse’s intrinsic *β*-cells, iPSC-derived *β*-cells from MODY3 patient (+/R200Q) were unable to fully compensate for the increased insulin requirement, and gradually developed phenotypes resembling those of *HNF1A* knockout *β*-cells, including a failure to secrete insulin in response to glucose, and reduced insulin to C-peptide secretion ratios. Correction of the R200Q mutation in MODY3 iPSC-derived *β*-cells protected cells from acquiring abnormal insulin secretion profiles. This study provides direct evidence that the *HNF1A* R200Q mutation is functionally consequential and pathogenic, leading to insulin secretion defects.

Our studies demonstrate the utility of stem cell-based models in defining the molecular physiology of *β*-cell failure in gene-specific diabetes in humans. Specifically, we show pleiotropic effects of *HNF1A* deficiency at several levels of *β*-cell biology and function, consistent with the diversity of its transcriptional targets. The clinical implications of these studies are several. We now have the technical capacity to create viable, functional human islet cell clusters capable of supporting glucose homeostasis in mice. β-cells generated by *in vitro* stem cell differentiation are now being tested for their ability to support glucose homeostasis in patients with autoimmune diabetes. In MODY patients, β-cell autoimmunity is not mechanistically involved, so that islets created from somatic cells of such patients should be immunologically tolerated by the patient. Using techniques described in this paper, the mutant alleles of MODY3 (or other) patients could be corrected in stem-cell derived islets. These cells could be used to restore glucose homeostasis in such subjects. Additionally, insights gained with regard to the molecular physiology of MODY3 may point to novel pharmacological interventions for those disorders, and possibly for more prevalent forms of diabetes.

## Material and methods

### Human subject and cell lines

Nine MODY3 subjects (Pt1 to Pt9), three MODY2 subjects (1068, 1133 and 1144) and four control subjects (1023, 1098, 1136 and 1015) were recruited at the Naomi Berrie Diabetes Center and monitored over 2-5 years. Samples were coded to protect subjects’ identity (**Fig. S2J**). Biopsies from MODY3 subjects (Pt1, Pt2 and Pt3) and one control subject (1023) were cut into small pieces (approximately 5x5 mm in size). 2-3 pieces of minced skin are placed next to a droplet of silicon in a well of six-well dish. A glass cover slip (22x22 mm) was placed over the biopsy pieces and silicon droplet. 5 ml of biopsy plating media was added and incubated for 5 days at 37°C. Biopsy pieces are then grown in culture medium for 3-4 weeks. Biopsy plating medium is composed of DMEM (Gibco, 10569), 10% FBS (GE Healthcare, SH30088.03HI), 1% GlutaMAX (Gibco, 35050061), 1% Anti-Anti (Gibco, 15240062), NEAA (Gibco, 11140-050), 0.1% 2-Mercaptoethanol (Gibco, 21985023) and nucleosides (Millipore ES-008-D). Culture medium contains DMEM (Gibco, 10569), 10% FBS (GE Healthcare, SH30088.03HI), 1% GlutaMAX (Gibco, 35050061) and 1% Pen-Strep (Gibco, 15070063). The hESCs (Mel1)(*21*) used in this manuscript is an NIH approved line. All research involving human subjects was approved by the Institutional Review Board of Columbia University Medical Center, and all participants provided written informed consent.

### Generation of iPSCs and cytogenic analysis of stem cells

Primary fibroblasts were reprogrammed into pluripotent stem cells using CytoTune™-iPS Sendai Reprogramming Kit (Invitrogen). 50,000 fibroblast cells (between passage 2-5) were seeded in a well of a six-well dish and allowed to recover overnight. The next day, cells were infected by Sendai virus expressing human transcription factors Oct4, Sox2, Klf4 and C-Myc mixed in fibroblast medium according to the manufacturer’s instructions. Two days later, the medium was exchanged for human ES medium supplemented with the ALK5 inhibitor SB431542 (2 µM; Stemgent), the MEK inhibitor PD0325901 (0.5 µM; Stemgent), and thiazovivin (0.5 µM; Stemgent). Human ES medium contained KO-DMEM (Gibco 10829), 15% KnockOut Serum Replacement (Gibco, 10828), 1% GlutMAX, 0.1% 2-Mercaptoethanol, 1% NEAA, 1% PenStrep and 0.1 ug/mL bFGF (all from Gibco). On day 7-10 post infection, cells are detached using TrypLE™ Express (Gibco, 12605036) and passaged onto mouse embryonic fibroblast feeder cells (GlobalStem CF-1 MEF IRR). Individual colonies of induced pluripotent stem cells were manually picked between day 21-28 post infection and each iPSC line was expanded from a single colony. All stem cells lines are cultured on human ES medium. Cytogenic analysis performed on 20 G-banded metaphase cells from each line by Cell Line Genetics Inc (**Fig. S2M**).

### Stem cell culture

hESC and iPSCs are grown on plates coated with primary mouse embryonic fibroblasts or MEFs (GlobalStem, CF-1 MEF IRR) and dissociated every 4-5 days using TrypLE™ Express (Life Technology, 12605036) for passaging. After dissociation, cells are suspended in human ES medium containing 10 µM ROCK inhibitor Y27632 (Selleckchem, S1049).

### *In silico* gRNA design

sgRNAs (**Table S1**) were designed using an online CRISPR design tool (crispr.mit.edu) and cloned into a gRNA cloning vector (Addgene, 41824) following option B from the gRNA Synthesis Protocol(*53*). The resulting vector is transformed into competent bacteria using Gibson Assembly® chemical transformation protocol (E5510). Single clones are picked and grown on 3 ml of LB broth for 16-18h at 37°C in a shaker (250 rpm); DNA is extracted and Sanger sequenced.

### CRISPR/Cas9 nucleofection and mutagenesis

Stem cells are cultured with human ES media containing ROCK inhibitor Y27632 (Selleckchem, S1049) 3h prior nucleofection, dissociated and filtered through a 70 µm cell strainer (Thermo Fisher Scientific, 8-771-2). Approximately 2x10^6^ cells are nucleofected (Lonza nucleofector, program A23) with 5 µg of Cas9-GFP plasmid (Addgene, 44719), 5 µg of sgRNA and 5 µg of ssDNA donor template (**Table S1**) using Human Stem Cell Nucleofector^TM^ kit 1 (Lonza, VVPH-5012) according to the manufactures protocol and cells replated. Cells are dissociated 48h later for GFP sorting using BD FACS Aria II cell sorter. As a quality control step, some unsorted cells were used to test the sgRNAs mutation efficiency using the Transgenomic SURVERYOR® mutation detection kit according to the manufactures protocol. Sorted cell are plated in a 10cm dish and single clones are picked 7-10 days post sorting, clones are further expanded and DNA extracted for PCR using *HNF1A* primers (**Table S1**). Amplicons are sent for Sanger sequencing to GENEWIZ and clones with indels are further validated by TOPO® TA cloning (Thermo Scientific, 450641) (at least 6 clones are picked) followed by Sanger sequencing. For hESC WT line, five different clonal lines were used for analysis throughout the study.

### Differentiation into pancreatic endocrine cells

Cells are grown to 80-90% confluency, dissociated and suspended in mTeSR™ medium (STEMCELL Technology, 05850) with 10 µM ROCK inhibitor Y27632 (Selleckchem, S1049) and plated in a 1:1 ratio into Matrigel-coated (Fisher Scientific, 354277) wells for differentiation. Differentiation was performed using a published protocol (*20*). The initial stages of differentiation were conducted in planar culture (d0-d11). For definitive endoderm stage (d1-d3) cells were cultured using STEMdiffTM Definitive Endoderm Differentiation Kit (Stemcell Technologies, 05110). For primitive gut stage (d4-d6), cells were cultured in RPMI containing GlutaMAX (Life Technology, 61870-127), 1% (v/v) Penicillin-Streptomycin (PS) (Thermo Fisher Scientific, 15070-063), 1% (v/v) B27 Serum-Free Supplement (50x) (Life Technology, 17504044) and 50 ng/ml FGF7 (R&D System, 251-KG). For posterior foregut stage (d7-d8), cells were cultured in DMEM containing GlutaMax, 1% (v/v) PS, 1% (v/v) B27, 0.25 µM KAAD-Cyclopamine (Stemgent, 04-0028), 2 µM Retinoic acid (Stemgent, 04-0021) and 0.25 µM LDN193189 (Stemgent, 04-0074). For pancreatic progenitor stage (d9-d11), cells were cultured in DMEM containing GlutaMax, 1% (v/v) PS, 1% (v/v) B27 and 50 ng/ml EGF (R&D System, 236-EG). Cells were then dissociated using TrypLE™ Express (Life Technology, 12605036) and seeded into low-attachment 96 well-plates (Corning, 7007) (1 well of 6 well-plate to 60 wells of 96-well-plate) for clustering step to form aggregates or clusters of endocrine cells in DMEM containing GlutaMax, 1% (v/v) PS, 1% (v/v) B27, 0.25 µM Cyclopamine, 1 µM thyroid hormone (T3) (Sigma, T6397), 10 µM Alk5i, 10 µM Zinc sulfate (Sigma-Aldrich, Z4750) and 10 µg/ml Heparin (Sigma-Aldrich, H3149) for 2 days (d12-d13). For pancreatic endocrine stage (d14-20) cells were cultured using DMEM containing GlutaMax, 1% (v/v) PS, 1% (v/v) B27, 100 nM LDN, 1 µM T3, 10 µM Alk5i, 10 µM Zinc sulfate, 10 µg/ml Heparin and 100 nM gamma-secretase inhibitor (DBZ) (EMD Millipore, 565789). For mature pancreatic endocrine stage (d21-d27) cells were cultured using DMEM containing GlutaMax, 1% (v/v) PS, 1% (v/v) B27, 1 µM T3, 10 µM Alk5i, 10 µM Zinc sulfate, 10 µg/ml Heparin, 1 mM N-acetyl cysteine (N-Cys) (Sigma-Aldrich, A9165-5G), 10 µM Trolox (EMD Millipore, 648471-500MG) and 2 µM R428 (Tyrosine kinase receptor AXL inhibitor) (ApexBio, A8329). From d1 to d11 media was changed every day and from d12 to d27 media was changed every other day. All differentiations were done for 27 to 30 days.

### *In vitro* insulin secretion and content

Static insulin secretion assay was performed at day 27-30 of differentiation; 10 islet-like clusters of cells were used per experiment. Islet-like clusters were pre-incubated for 1 hour in Kreb’s Ringer Buffer (128mM NACL, 5mM KCL, 2.7mM CaCl2, 1.2mM MgSO4, 1mM NaHPO4, 1.2mM KH2PO4, 5mM NaHCO3, 10mM HEPES, 0.1% Bovine Serum Albumin, pH=7.4) containing 3.3 mM glucose, washed and incubated for another hour in 3.3 mM glucose and the medium was collected. Subsequently, 200 µl of buffer containing 16.7 mM glucose or 400 µM tolbutamide (abcam, ab120278) or 30.5 mM KCl was used to treat cells for 1 hour, after which the medium was collected. Insulin content was measured by acid ethanol extraction; cells are resuspended in 50 µl of water and sonicated for 15 seconds. The sonicate is mixed with acid ethanol (0.18 M HCl in 96% ethanol (vol/vol)), in a 1:3 ratio of sonicate to acid ethanol. The mixed solution is incubated at 4°C for 12 hours. Human C-peptide, human insulin and human proinsulin secretion and content was measured using Ultrasensitive C-peptide ELISA (Mercodia, 10-1141-01), Insulin ELISA (Mercodia, 10-1113-01) and Proinsulin ELISA (Mercodia, 10-1118-01) kit according to the manufactures protocol. All samples were handled the same way.

### Dynamic insulin secretion

Perifusion was performed as described previously(*54*), 20 randomly chosen cluster of cells were examined using a Biorep Technologies (Miami, FL) perifusion system. Clusters were perifused with Kreb’s buffer [115mM NaCl, 5mM KCl, 24mM NaHCO_3_, 1mM MgCl_2_, 2.2mM CaCl_2_ at pH 7.4] supplemented with 0.17% bovine serum albumin and 3.3 mM glucose (26 min), followed by 16.7 mM glucose (35 min), 3.3 mM glucose (15 min), 400 µM tolbutamide in 3.3 mM glucose (35 min), 20 mM KCl plus 3.3 mM glucose (10 min) and finally 3.3 mM glucose (15 min). Medium was collected at a flow rate 100 µl/min to assess insulin secretion. Insulin concentration was measured using an Insulin ELISA kit (Alpco, 80-INSHU-E01.1). Cluster of cells were collected at the end of the study and placed in acidified ethanol overnight to determine total insulin levels. All samples were handled the same way.

### Calcium Imaging

Stem cell-derived β-cells were dissociated from clusters and plated onto 35 mm glass-bottom dishes coated with 5% Matrigel (Fisher Scientific, 354277). Cells were washed twice with basal KRBH solution composed of (mM): 128 NACl, 5 KCl, 2.7 CaCl_2_, 1.2 MgSO4, 1 NaHPO_4_, 1.2 KH_2_PO_4_, 5 NaHCO_3_, 10 HEPES, 0.1% Bovine Serum Albumin, 3.3 glucose, pH to 7.4. Cells were then incubated in the same solution containing 1 uM fura-2, AM (ThermoFischer, F1221) with 0.05% pluronic F-127 in DMSO (ThermoFischer, P3000MP) for 15 min. at 37°C, 5% CO_2_. Cells were washed twice with basal KRBH solution and then imaged on an inverted Nikon Ti-eclipse microscope with a Nikon Plan fluor 20x objective (0.45 N.A.). Fura-2 measurements were collected at excitation wavelengths of 340 and 380 nm using EasyRatioPro (HORIBA Scientific). Stimulation solutions included either 16.7 mM glucose, 600 µM tolbutamide, or 30.5 mM KCl, with NaCl concentrations adjusted accordingly to balance osmolarity with KRBH solution. Calcium concentrations were calculated as follows:

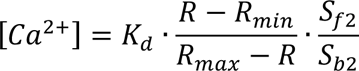

where K_d_, the apparent Ca^2+^ binding affinity to Fura-2, was assumed to be 225 nM and *R*_max_, *R*_min_, *S*_f2_, and *S*_b2_ values were obtained using 10 µm ionomycin with either no Ca^2+^ or 2 mM Ca^2+^, values (*55, 56*).

### Immunohistochemistry

Clusters of cells and tissue (graft or pancreas) were fixed with 4% paraformaldehyde for 20 minutes at room temperature or 24h at 4°C respectively, washed in PBS, then dehydrated with 30% sucrose for 24 hours. Cells and tissues were washed in PBS, then cryopreserved in frozen OCT and sectioned at 7 µm on microscope slides. Slides are then incubated with blocking solution containing 3% donkey serum with 0.1% triton X-100 diluted in PBS for 1h at room temperature. Primary antibodies (**Table S7**) are added in block solution overnight at 4°C, washed three times for 5 minutes in PBST (PBS with 0.1% Tween-20) and incubated with secondary antibodies (**Table S7**) diluted in block solution for 1h at room temperature, washed three times in PBST containing 10 µg/ml DAPI for 5 minutes and mounted with a coverslip in fluorescent mounting media (Dako, S3023). All images were taken using a Zeiss LSM 710 confocal microscope and quantified manually using ImageJ software. Quantification of cell type at the definitive and pancreatic progenitor stage (planar culture) were done by randomly choosing 4 sections in the well and averaging the percentage of cells from all sections. For cluster of cells (3D culture) and tissue (graft or pancreas), they were cut in 3-6 sections spaced 20 µm and 100 µm respectively and cell type was quantified in each section as a percentage. Final quantification result is the average of all sections. Each cluster of cell is comprised of ∼10’000 cells.

### Flow Cytometry

Clusters of endocrine cells are dissociated into single cells using TrypLE™ Express (Life Technology, 12605036). Cells are then fixed with 4% paraformaldehyde for 20 minutes at room temperature followed by 10 minutes permeabilization with cold methanol at -20°C. Cells are washed with 3% donkey serum diluted in PBS and primary antibodies diluted (**Table S7**) in blocking solution containing 3% donkey serum with 0.1% triton X-100 diluted in PBS overnight at 4°C. For tunel assay (Biotium, 30074) cells are incubated with fluorescent biotinylated nucleotide conjugate according to the manufacturer’s instructions. Cells are washed in PBST (PBS with 0.1% Tween-20) and incubated with secondary antibodies diluted (**Table S7**) in block solution for 1h at room temperature. Cells are washed and filtered with a BD Falcon 12×75–mm tube with a cell strainer cap (BD Biosciences, 352235) and analyzed by flow cytometer. Data were analyzed using FlowJo software. Gating for flow cytometry was determined by incubating the cells only with secondary antibodies (negative control) and was consistent across experiments (shown in magenta in figures).

### Western Blot

Cluster of endocrine cells are dissociated into single cells using TrypLE™ Express (Life Technology, 12605036). Cells are lysed using RIPA buffer (NP40 1%, NaCl 150mM, EDTA 1mM, Tris-HCl pH 7.5 50nM, SDS 0.1%, Sodium Deoxychol 0.5%, NaF 10mM, protease inhibitor tablet) and whole-cell lysates obtained by subsequent centrifugation. Immunoblots were incubated with primary antibodies against HNF1A (abcam, ab96777, 1:500) and β-Tubulin-III (Sigma, T2200, 1:500) (**Table S7**).

### Bulk RNA sequencing

Around 200 clusters of cells are dissociated and sorted for GFP using BD FACS Aria II cell sorter. Sorted cells are pelleted by centrifugation and RNA extracted using total the RNA purification micro (Norgen Biotek, 35300) kit according to the manufactures protocol. RNA quality and concentration is determined by an Agilent bioanalyzer and RNAseq performed by the Columbia University Genome Center. A poly-A pull-down was used to enrich mRNAs from total RNA samples (200ng-1ug per sample, RIN>8 required) and proceed on library preparation by using Illumina TruSeq RNA prep kit. Libraries are then sequenced using an Illumina HiSeq2000. RTA (Illumina) was used for base calling and bcl2fastq (version 1.8.4) for converting BCL to fastq format, coupled with adaptor trimming. The reads were mapped to a reference genome (Human: NCBI/build37.2; Mouse: UCSC/mm9) using Tophat(*57*) (version 2.1.0) with 4 mismatches (--read-mismatches = 4) and 10 maximum multiple hits (--max-multihits = 10). To tackle the mapping issue of reads that are from exon-exon junctions, Tophat infers novel exon-exon junctions ab initio, and combine them with junctions from known mRNA sequences (refgenes) as the reference annotation. The relative abundance (aka expression level) of genes and splice isoforms is estimated using cufflinks(*58*) (version 2.0.2) with default settings. Differentially expressed genes are tested under various conditions using DEseq(*59*). An R package based on a negative binomial distribution that models the number reads from RNA-seq experiments to test for differential expression.

### Single-cell RNA sequencing

Around 100 clusters of cells per genotype at day 27 of differentiation are dissociated and sorted for GFP using BD FACS Aria II cell sorter. Single cell RNA-Seq data from total of 271 hESC-derived cells (113 for WT and 158 for KO) were obtained. Raw reads were aligned to the hg19 reference genome using STAR (2.5.2b) and gene quantification were performed using FeatureCounts (1.5.2). Quality control was conducted as follows: cells with low mapped reads (<1,000,000), low exon count (<100,000), low exon mapped rate (<0.2), high intergenic mapped rate (>0.3), low detected gene count (<1,000) and with more than 10% Mitochondria genes’ count were removed. In addition, genes with <10 non-zero counts across all the cells were eliminated. All the downstream analysis was done with in R-3.4.1. Filtered raw count data were normalized and scaled by Seurat package and only variable genes generated by Seurat were used to do PCA and t-SNE. n=3 for each genotype.

### 10X genomics single-cell RNA sequencing

Around 100 clusters of cells per genotype at day 27 of differentiation are dissociated. Single cells were suspended in PBS+0.04% BSA. Cellular suspensions (∼6,000 cells) were loaded on a Chromium Single Cell Instrument (10X Genomics) to generate single cell GEMs. Single-cell RNAseq libraries were prepared using Chromium Single cell 3¢ Library, Gel beads & Mutiplex kit (10X Genomics). Sequencing was performed on Illumina NextSeq500 using the following read length: 59-bp Read1 for transcript read, 14-bp I7 Index for Cell Barcode read, 8-bp I5 Index for sample index read, and 10-bp Read2 for UMI read. The samples were sequenced in ten different libraries (batches). Among all these samples, Batch 980379 and Batch 993091 (2 libraries) are HNF1A mutants (R200Q/R200Q), Batch 980380 and Batch 993090 (2 libraries) are HNF1A knockouts (KO) while Batch 989516 and Batch 993089 (2 libraries) are HNF1A wildtype cells (WT). In addition, to increase the sequencing depth, each batch was sequenced from 7 flowcells. The raw fastq files was processed via 10X Cell Ranger pipeline (cellranger-2.1.1). To note that GRCh38 reference genome was applied for the whole analysis. “cellranger count” with “--expect-cells=3,000” was run for each library respectively and “cellranger aggr” with default parameters was run to aggregate results generated from multiple libraries. A normalized umi-count matrix for all the cells was then generated. As for the quality control, cell and gene filtering process, cells with umi-count<5,000 and umi-count>40,000 and cells with <2,000 detected genes were removed. Also, genes with <∼1% (25) non-zero counts across all the cells were eliminated from the analysis. To get rid of the Mitochondria cells, cells with more than 10% Mitochondria genes’ count were also removed. All the downstream analysis was done with Seurat package in R-3.4.1. Every batch is an independent differentiation (biological replicates). n=2 for each genotype.

### Accession numbers

Single cell RNA sequencing and bulk RNA sequencing data were deposited in NCBI’s Gene Expression Omnibus (GEO) database and accession number is GSE128331 and GSE129653.

### Cell transplantation and *in vivo* assays

After 27-30 days of differentiation, 50 µg of stem cell-derived pancreatic cell clusters (∼180 islet-like clusters of 300-350 µm diameter; each cluster comprised ∼10,000 cells in total) including all islet endocrine cell types were suspended in 30 µl of ice-cold Matrigel (Fisher Scientific, 354277) and loaded into ice-cold 1ml syringe with a 21G needle for transplantation. 6-10 weeks old male immunodeficient NSG (NOD.Cg-Prkdcscid Il2rgtm1Wjl/SzJ, Stock No: 005557, The Jackson Laboratories) mice were transplanted in the ventral and medial muscles (medial thigh and posterior lower) of the left thigh. Every 2 weeks post-transplantation, human C-peptide and human insulin was determined in the plasma of recipient mice at fed state (morning) for 30 weeks. An intraperitoneal glucose tolerance test was performed 18-24 weeks (before STZ injection) and 28-30 weeks (after STZ injection) post-transplantation by injecting 2 g/kg body weight of 20% D-glucose (Sigma, G8270) in PBS after overnight fasting (16-18h). For glibenclamide (Sigma, G0639) administration, 1mg/kg body weight of the drug diluted in PBS was i.p injected 24-28 weeks (before STZ injection) post-transplantation, all mice were at fed state but food was retrieved during the experiment. Blood was collected by tail bleeding in heparinized Eppendorf tubes at time points; plasma was isolated by centrifugation for 15 minutes at 2000g at 4°C. Human C-peptide was measured using Mercodia Ultrasensitive C-peptide ELISA (10-1141-01) and human insulin using Mercodia Insulin ELISA (10-1113-01) kit according to the manufactures protocol. All samples were handled the same way. Insulin detection limit is 1mU/L as determined by the methodology described in the kit. Blood glucose levels were measured using a glucometer (FreeStyle Lite) and HbA1C using Siemens DCA 2000 Vantage Reagent kit (Siemens, 5035C). To induce diabetes, mice were i.p injected for 5 consecutive days with 40 mg/kg body weight of Streptozotocin (STZ) (Sigma-Aldrich, S0130-1G) in PBS 24-28 weeks post-transplantation. All experiment procedures were performed according to Columbia University approved IACUC protocols.

### Bioluminescence and fluorescence imaging

NSG mice transplanted with *GAPDH^Luciferase/wt^* and *INS^GFP/wt^* double reporter hESCs lines were i.p injected with 150mg/kg body weight of D-luciferin potassium salt (Gold Biotechnology, luck-2G) in PBS 15 minutes before imaging on a IVIS spectrum optical imaging system (PerkinElmer). Signals were acquired with 1-minute exposure and analyzed using the Living image analysis software (Xenogen Corp.). Circular regions of interest (ROI) of the same size for all experiments were drawn around the signal in the left thigh and photons emitted over the time of exposure within the ROI measured. Bioluminescence was measured every 2 weeks post-transplantation for 30 weeks and after isolation of the graft at that time point. Luminescence was measured as described for bioluminescence after isolation of the graft. Background signals were subtracted from a nearby region and were consistent over time.

### Electron microscopy

Fixed cells/tissues were incubated with 2.5% glutaraldehyde and 2% paraformaldehyde in 100 mM sodium cacodylate buffer (pH 7.4) overnight. Samples were then treated with 1% osmium tetroxide in 100 mM sodium cacodylate buffer for 1 h, washed in distilled water four times (10 min/wash), and then treated with 2% aqueous uranyl acetate overnight at 4°C in the dark. Samples were washed and sequentially dehydrated with increasing concentrations of acetone (20, 30, 50, 70, 90, and 100%) for 30 min each, followed by three additional treatments with 100% acetone for 20 min each. Samples were then infiltrated with increasing concentrations of Spurr’s resin (25% for 1 h, 50% for 1 h, 75% for 1 h, 100% for 1 h, 100% overnight at room temperature), and then incubated overnight at 70°C in a resin mold. Sections of 50–90 nm were cut on a Leica ultramicrotome with a diamond knife. Imaging was performed on an FEI Talos L120C operating at 120kV.

### Immunogold electron microscopy

70nm thick sections from the embedded samples were placed on to carbon formvar 75 mesh nickel grids and etched using 4% sodium metaperidotate for 10 minutes before being washed twice in distilled water and then blocked for 1 hour. Grids were incubated with the primary C-peptide antibody (**Table S7**) (1 in 10 dilution) for 1 hour at 4C overnight. Next day grids underwent seven washes in 1xPBS and then incubated in anti-rat 6nm gold secondary (1 in 50 dilution) for 1 hour (**Table S7**). After this the grid was washed seven times in 1xPBS and twice in distilled water. Samples were then imaged on a ThermoFisher Talos L120C operating at 120kV.

### Statistical analysis

All statistics were performed using Prism GraphPad software (La Jolla, CA). Normality of data set distributions was assessed by Shapiro-Wilk test and D’Agostino & Pearson omnibus tests. Normally distributed data were analyzed by unpaired two-tailed *t-*test. For data not normally distributed we used a two-tailed Mann-Whitney test. Data are expressed as mean ± standard error of the mean (SEM). *P*<0.05 was considered statistically significant. *p<0.05, **p<0.01, ***p<0.001. n.s indicates a non-significant difference.

## Supplementary materials

Fig. S1. Schematic overview of main figures.

Fig. S2. Generation of isogenic cell lines with *HNF1A* mutations in hESCs and MODY3 iPSCs.

Fig. S3. *HNF1A* is not required to generate pancreatic endocrine cells *in vitro*.

Fig. S4. Identification of thirteen stem cell-derived cell populations by single cell RNAseq *in vitro*.

Fig. S5. Identification of scβ- and sc*α*-like-cells and their gene regulatory network *in vitro*. **Fig. S6**. *HNF1A* deficiency results in a bias of endocrine cells towards the *α*-cell fate *in vitro.* **Fig. S7**. *HNF1A* deficiency results in a bias of endocrine cells towards the *α*-cell fate *in vivo*.

Fig. S8. *HNF1A* deficiency affects insulin secretion in association with *CACNA1A* and *SYT13* down-regulation *in vitro*.

Fig. S9. *HNF1A* deficiency causes abnormal insulin granule structure.

Fig. S10. *HNF1A* deficient scβ-cells are unable to maintain glucose homeostasis in diabetic mice. **Fig. S11**. *HNF1A* R200Q mutation is pathogenic and causes developmental bias towards the *α*-cell fate *in vitro*.

Fig. S12. MODY3 iPSC-derived β-cells are initially glucose responsive.

Fig. S13. *HNF1A* haploinsufficiency gradually impairs scβ-cell function *in vivo*.

## Supplementary tables

Table S1. Primer and oligo (sgRNA & ssDNA) sequence used for qPCR and CRISPR/Cas9 with *HNF1A* sgRNA off-target sites characterization.

Table S2. Sanger sequencing for *HNF1A* sgRNAs off-target genes in different cell lines.

Table S3. List of down/up-regulated genes from bulk RNAseq transcriptome of *INS^GFP/wt^* sorted cells *in vitro*.

Table S4. GOTERM and KEGG analysis from single cell RNAseq transcriptome of *INS^GFP/wt^*sorted cells *in vitro*.

Table S5. GOTERM and KEGG analysis from single cell RNAseq transcriptome of unsorted SC-islet-like cells *in vitro*.

Table S6. Genotypes and clinical phenotypes of MODY3 patients. Anthropometry and clinical measurements of non-fasting plasma insulin (pM), C-peptide (pM), INS(pM)/C-PEP(pM) ratio, glucagon (pM) and proinsulin (pM) from MODY3 subjects (n=9), MODY2 subjects (n=3) and non-diabetic subjects (n=4). <1 indicates below detection limit for human insulin by ELISA kit. Red indicates low insulin to C-peptide ratio. MODY3 patient 2 (Pt2), mother of patient 3 (Pt3) and segregating for same *HNF1A* (+/R200Q) mutation had a history of gestational diabetes in 2 pregnancies and was diagnosed as having T2D in 2003 at age 38. She was managed with oral agents until 2015 for increasing hyperglycemia (292 mg/dl and HbA1c of 10.3%) when insulin was initiated and sulfonylurea discontinued. Since her last visit in 2016, fasting glucose levels remained elevated (216 mg/dl and HbA1c of 11.9%). MODY3 patient 3 (Pt3) was misdiagnosed as having T1D at age 13 and was found to have a mutation in *HNF1A* (+/R200Q) at age 15. Insulin was discontinued and sulfonylurea treatment prescribed. Fasting glucose levels at age 19 were 177 mg/dl with an HbA1C of 7.8%, for which the sulfonylurea dosage was increased. Since then, Pt3’s fasting glucose and HbA1C levels have risen progressively, reaching 338 mg/dl and 6.9% by 2017 at age 22. During the patient’s last visit in 2017, non-fasting C-peptide levels were within the normal range (724 pM) but insulin was undetectable (0 pM).

Table S7. Primary and secondary antibodies used for immunohistochemistry

## Acknowledgments

We would like to thank Haiqing Hua for helping to generate the MODY3 patient iPSC lines and Giacomo Diedenhofen for helping with western blots for HNF1A.

## Author Contributions

[B.J.G] designed the studies, performed differentiation, transplantation of all cell lines and downstream experiments, gathered the data, analyzed the data and wrote the paper with input from all authors. [D.E and R.L] designed the studies, discussed results, reviewed and edited the manuscript. [D.E] performed skin biopsy stem cell derivation and [B.J.G] plasma isolation from subjects. [B.J.G, H.Z, J.L and Y.S] designed, analyzed and compiled the single-cell RNA sequencing data. [B.J.G and C.N.G] performed EMC imaging. [W.K.C] provided with genetic sequence information of MODY3 subjects. [Y.X, Y.W and J.O] performed microfluidic perfusion and generated data. [J.N, D.J.W and H.M.C] performed calcium imaging, measurements and data interpretation. [J.G] performed single-cell RNA sequencing. [R.S.G] performed oversight of human subject research and provided de-identified clinical information. [D.E, C.A.L and R.L] provided guidance in the design and interpretation of studies.

---

**Fig. S1.**
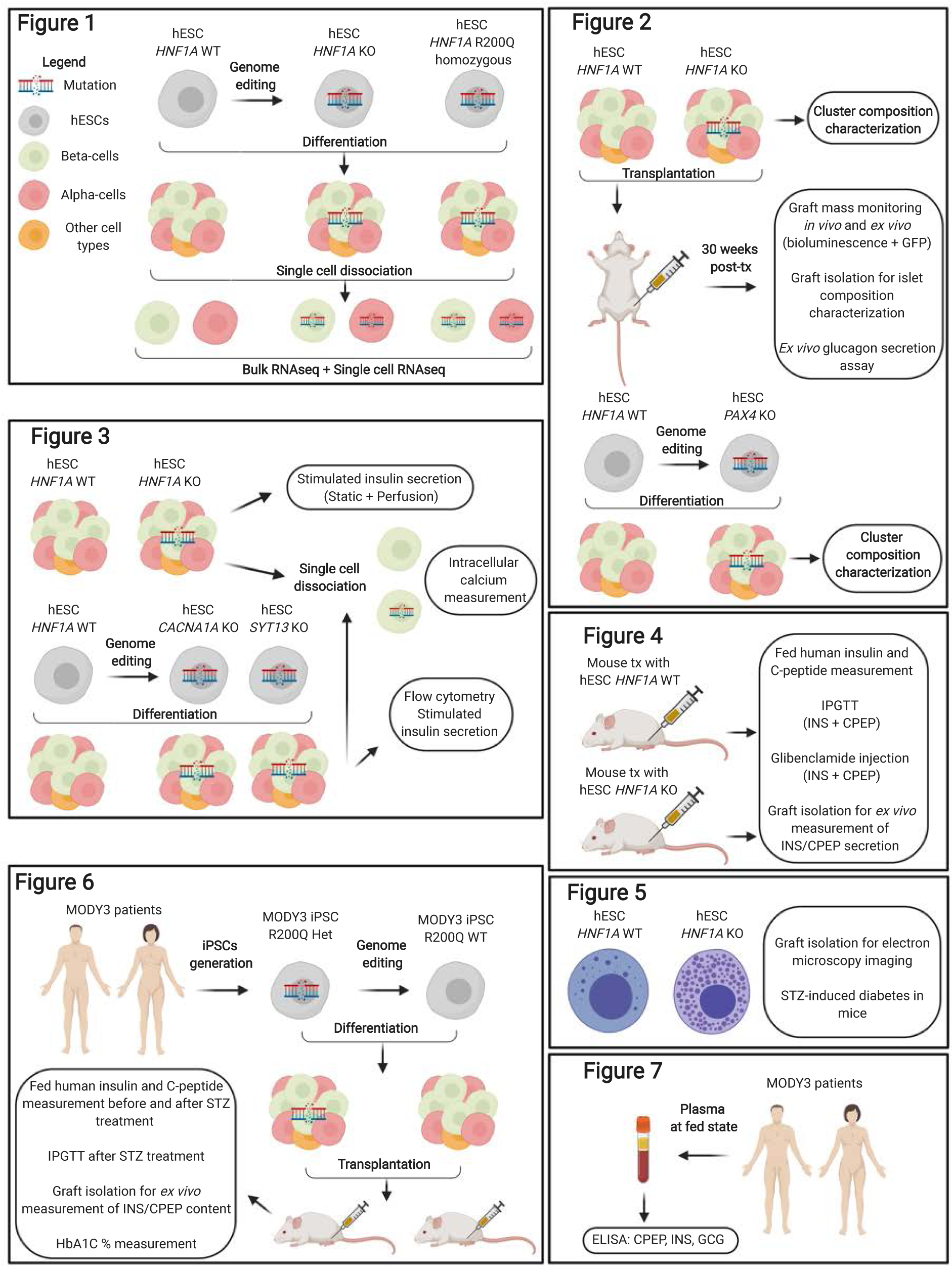
Schematic overview of main figures. IPGTT = intraperitoneal glucose tolerance test, STZ = streptozotocin, IHC = immunohistochemistry, MODY = maturity onset diabetes of the young, Het = heterozygous, WT = wild type. Designed using Biorender.

**Fig. S2.**
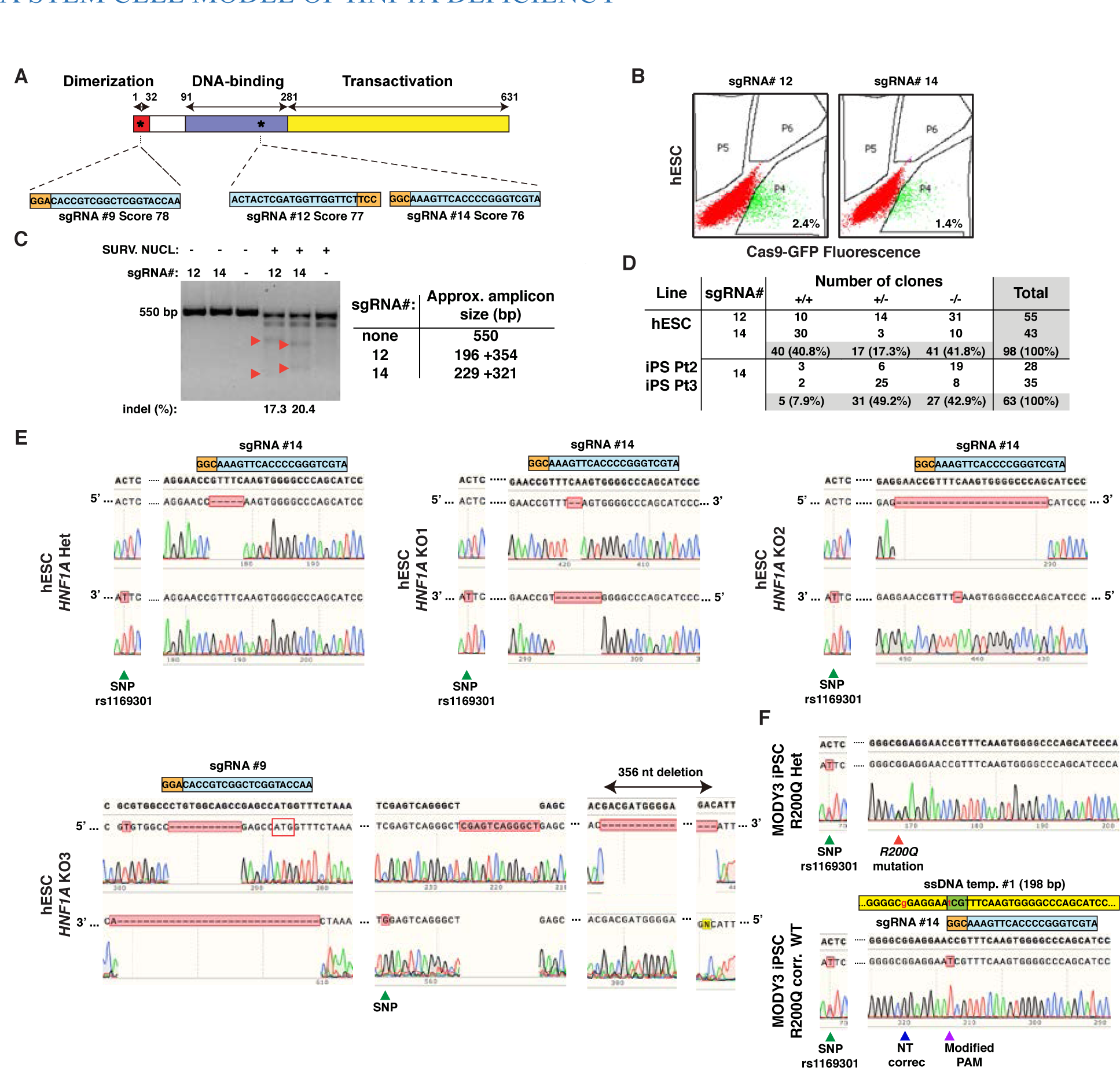
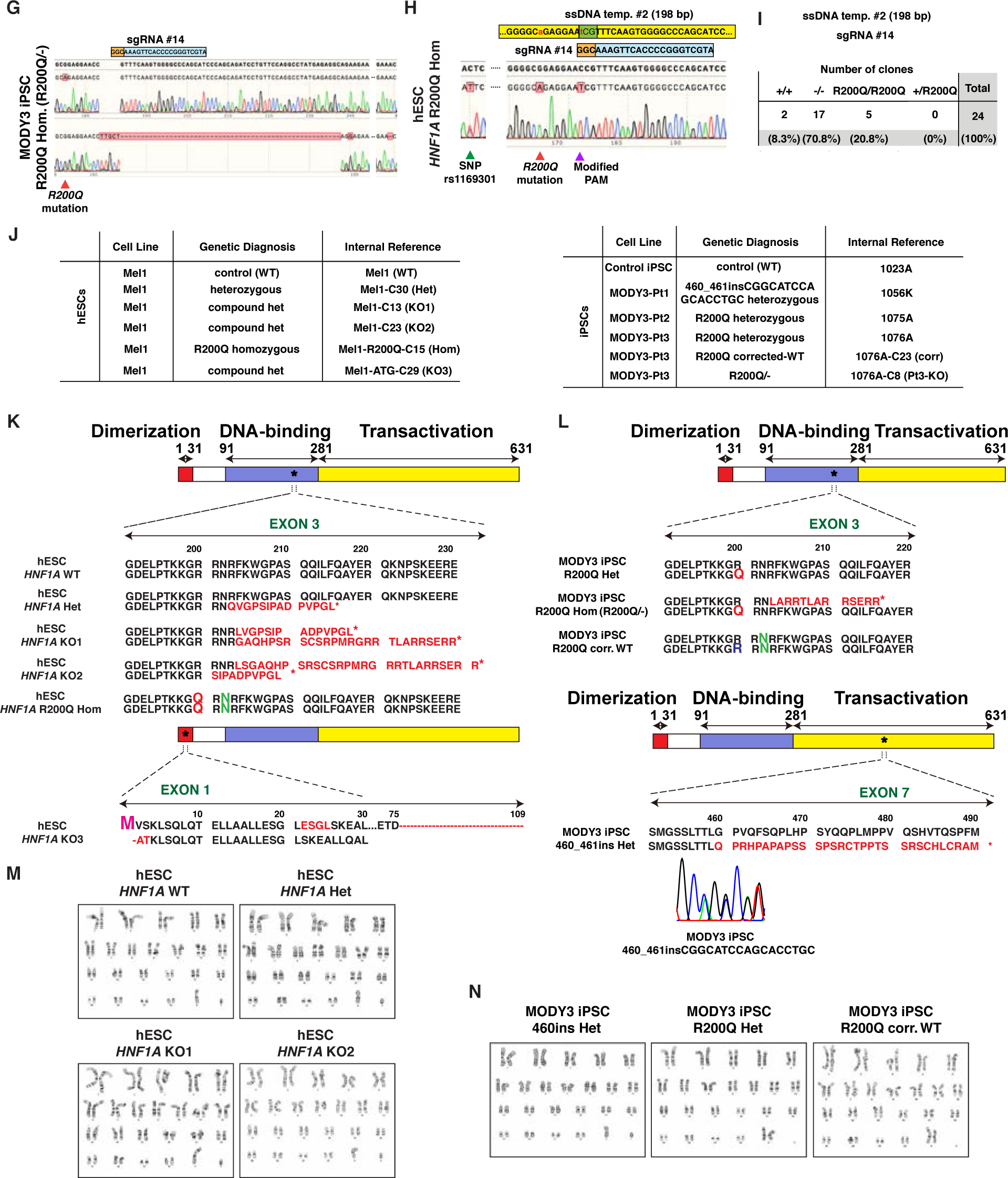
Generation of isogenic cell lines with *HNF1A* mutations in hESCs and MODY3 iPSCs. **(A)** Diagram illustrating functional domains of HNF1A protein with sgRNAs #12 (ontarget score 77) and #14 (on-target score 76) targeting the DNA-binding domain (exon 3). **(B)** FACs sorting of hESCs nucleofected with Cas9-GFP and sgRNA#12 or #14 with 2.4% and 1.4% trasnfection efficiency. **(C)** Surveyor assay to assess sgRNA indel efficiency in *HNF1A* gene. *HNF1A* PCR products with (+) or without (-) surveyor nuclease or sgRNA indicating cutting efficiency of 17.3% and 20.4% for sgRNA#12 and sgRNA#14, respectively, with indicated amplicon sizes (Table). The surveyor assay cleaves DNA at the site of heterozygosity. Bands with amplicon size of ∼500bp are due to the linked heterozygous SNPs rs1169301, creating cleavage. **(D)** Table showing the percentage of mutant lines generated after CRISPR/Cas9 in hESCs leading to 17.3% heterozygous clones (17 clones out of 98 total) and 41.8% compound heterozygous clones (41 clones out of 98 total), indicating a 59.2% of mutation efficiency (58 clones out of 98 total). 4 clones of the 98 (4%) appeared homozygous by Sanger sequencing, but also lacked two linked SNPs, rs1169301 (102 bp from PAM sequence of sgRNAs#14 to SNP) and rs2071190 (130 bp from PAM sequence of sgRNAs#14 to SNP), which are both located in the second intron. These homozygous lines (4/98, 4%) could be caused by a large deletion, and were not used for further studies. In MODY3 iPSC, 92.1% of clones were (+/-, heterozygous) and (-/-, compound heterozygous). Mutation-correction efficiency of MODY3 iPSC Pt2/Pt3 was 7.9% (+/+, wild type) from total number of clones. **(E)** Sanger sequencing results showing heterozygous (Het) and knockout frameshift mutations (KO1 and KO2) from both alleles in exon3 and knockout frameshift mutations (KO3) in exon1 near the start codon (red box). **(F)** Sanger sequencing showing heterozygous R200Q mutation (red arrow) in MODY3 iPSC line, after correction of the mutation (blue arrow) in MODY3 iPSC line (MODY3 iPSC R200Q-corrected WT) using a ssDNA template donor #1 (198 bp). **(G)** Sanger sequencing after introduction of a frameshift mutations in the wild type allele of MODY3 iPSC line (R200Q/-). **(H)** Sanger sequencing results showing R200Q homozygous mutation introduction (red arrow) in hESCs using a ssDNA template donor #2 (198 bp). PAM sequence in template donor was modified (C>T, silent mutation) to increase recombination efficiency (purple arrow). Heterozygosity at the upstream rs1169301 SNP C>T in *HNF1A* (green arrow; 102 bp from PAM sequence) indicates that both alleles are detected. All Sanger sequencing results were verified by TOPO® cloning of at least six clones per genotype. **(I)** Table showing the percentage of clones with R200Q homozygous mutation generated (20.8%) in hESCs from total number of clones. **(J)** Genetic information of cell lines (hESCs and iPSCs) included in the study**. (K)** Frameshift mutations in exon 3 of hESC cell lines leading to premature stop-codons (*) and generation of truncated versions of HNF1A protein (different amino acids from WT in red). For *HNF1A* KO3, mutation is at start codon in exon 1, indicated in red are different amino acids from WT sequence, red dash are deletions and in magenta the start codon. **(L)** Mutations in exon 3 of iPSC cell lines leading to heterozygous amino acid substitution (R200Q heterozygous), generation of R200Q homozygous (R200Q/-) and isogenic R200Q-corrected WT lines (+/+). For MODY3 iPSC 460_461ins Het line, CGGCATCCAGCACCTGC insertion is in exon 7 as shown by Sanger Sequencing, leading to frameshift mutations and premature stop codon (*). Different amino acids from WT in red, amino acid correction in bleu and amino acid silent mutation in green. **(M)** Cytogenic analysis performed on 20 G-banded metaphase cells from hESC and iPSC lines indicating normal karyotypes.

**Fig. S3.**
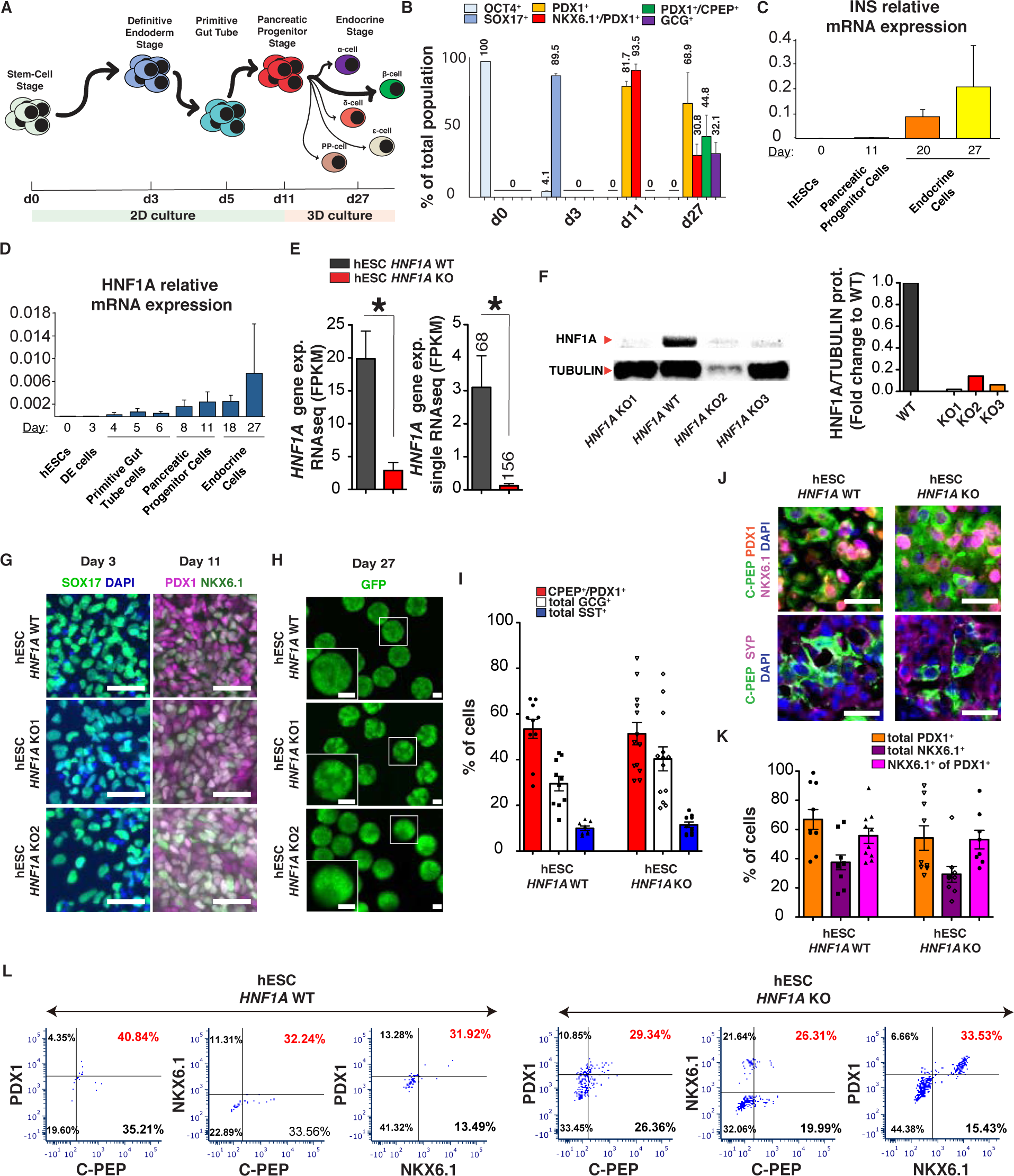
*HNF1A* is not required to generate pancreatic endocrine cells *in vitro*. **(A)** Overview of the different stages of stem cell differentiation to pancreatic progenitor cells. The initial stages of differentiation were conducted in planar culture from day 0 to day 11 followed by 3D clustering on day 12 to form aggregates or clusters of endocrine cells on day 27. **(B)** Percentage of total cell population by immunohistochemistry (IHC) expressing key markers of pluripotency stage (OCT4, d0), definitive endoderm stage (SOX17, d3), pancreatic progenitor stage (PDX1 and NKX6.1, d11) and endocrine stage (CPEP, c-peptide; GCG, glucagon, d27). **(C)** *INS* and **(D)** *HNF1A* relative mRNA expression levels determined by qPCR throughout the differentiation from hESC *HNF1A* WT cells (n=3-7). Each independent biological replicate (n) consists of 2-3 technical replicates for all experiments. **(E)** *HNF1A* gene expression levels (FPKM) determined by RNAseq after FACs sorting of *INS^GFP/wt^* positive cells from hESC-derived endocrine cells (WT n=3 and KO n=3) on day 27 of differentiation *in vitro*. Numbers on top of histogram denotes total number of single cells used for analysis. **(F)** Western blot for HNF1A (67 kDa) and β-Tubulin-III (55 kDa) from hESC-derived endocrine cells with respective quantification on day 27 of differentiation *in vitro*. **(G)** Representative IHC staining of hESC-derived cells *in vitro* at definitive endoderm stage with SOX17 (day 3) and pancreatic progenitor stage with PDX1 and NKX6.1 (day 11). Scale bars: 50 µm. White cells are PDX1/NKX6.1 double positive cells. **(H)** Representative GFP images. Scale bars at 200 µm. **(I)** Percentage of scβ-like-cells (CPEP^+^/PDX1^+^ in red), sc*α*-like-cells (GCG^+^ in white) and scδ-like-cells (SST^+^ in blue) by IHC (day 27). **(J)** Representative IHC image of hESC-derived endocrine cell lines with for indicated markers with **(K)** quantification of PDX1^+^ and NKX6.1^+^ cells (day 27). (**L**) CPEP^+^, PDX1^+^ and NKX6.1^+^ populations by flow cytometry at day 27 of differentiation *in vitro*. Scale bars: 20 µm. 20 clusters (∼10k cells per cluster) of endocrine cells were used flow cytometry. For scatter plots, each point in plots represents an independent biological experiment (n). Data are represented as mean ± SEM. p-values: *p<0.05, **p<0.01, ***p<0.001, two-tailed t-test.

**Fig. S4.**
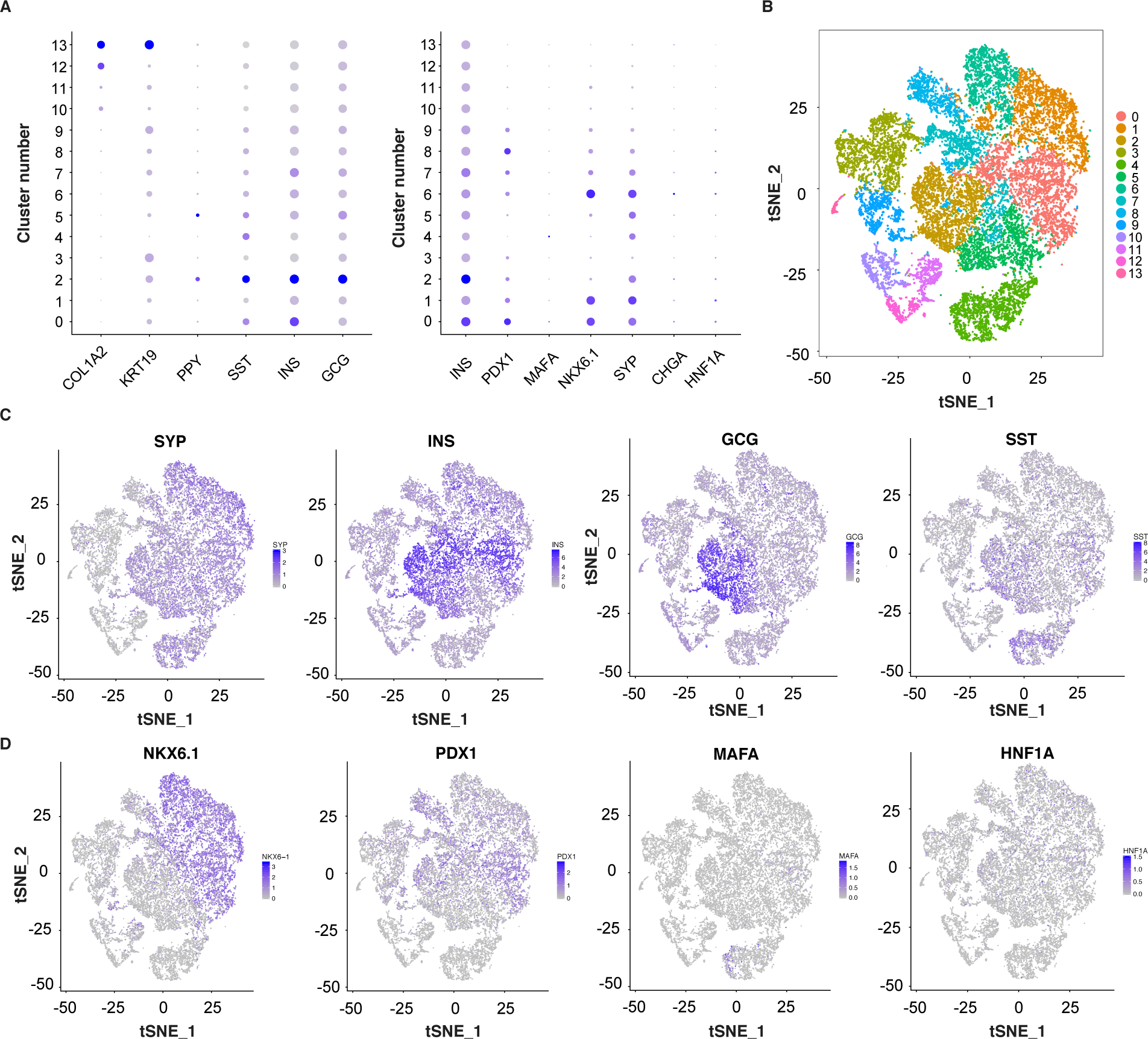
Identification of thirteen stem cell-derived cell populations by single cell RNAseq *in vitro*. **(A-D)** Single cell RNA sequencing of 22164 (all genotypes combined) unsorted hESC-derived endocrine cells (n=3 for each genotype) *in vitro*. **(A)** Cell type clustering of cells based on their normalized expression of different pancreatic markers: *COL1A2* (pancreatic stellate cell), *KRT19* (ductal cell), *PPY* (γ-cell), *SST* (δ-cell), *INS* (β-cell), and *GCG* (α-cell). A total of 13 clusters were identified. Size is the percentage of cells expressing the indicated marker and color intensity the average expression level of the indicated marker, where blue indicates the highest expression level and gray the lowest. **(B)** Feature plot based on tSNE projection of cells where the colors denote different cell type clusters identified (total of 13 identified) by *HNF1A* genotype line via Louvain algorithm performed by Seurat. **(C)** Feature plot based on tSNE projection of cells based on the normalized expression of endocrine markers *SYP* (synaptophysin), *INS* (insulin), *GCG* (glucagon) and *SST* (somatostatin) and **(D)** pancreatic transcription factors (*NKX6.1*, *PDX1*, *MAFA* and *HNF1A*) where blue indicates the highest expression level and gray the lowest. Genes expressed at low levels will have fewer labeled cells because of dropout. All stem cell differentiations were done for 27-30 days.

**Fig. S5.**
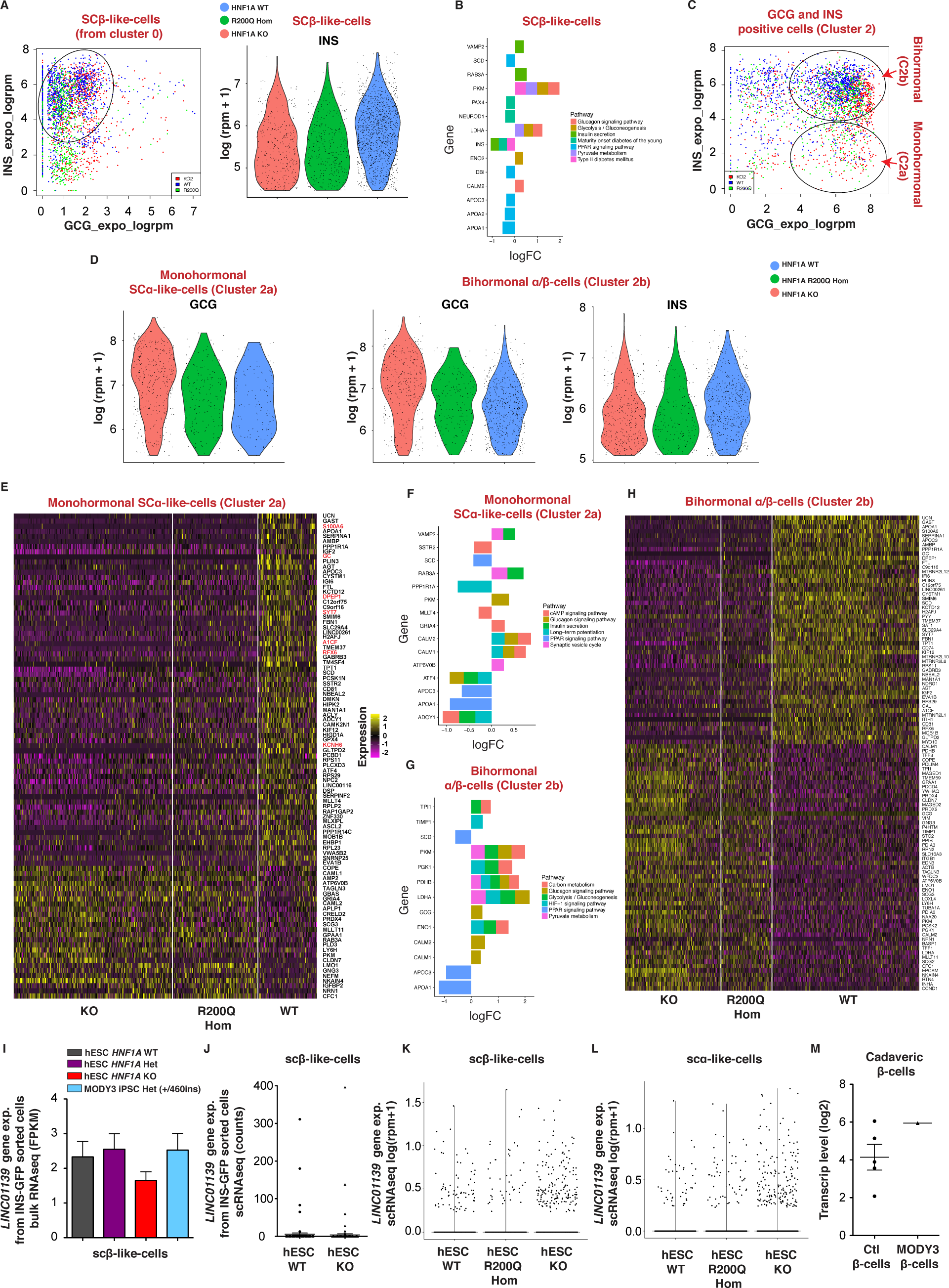
Identification of scβ- and sc*α*-like-cells and their gene regulatory network *in vitro*. **(A)** Scatter plot of 3133 hESC-derived β-like-cells from cluster 0 displayed (**Fig. S4**) based on *GCG* and *INS* expression (expo_logrpm). Total cell number from all genotypes combined. Encircled are scβ-like-cells (total of 1846 cells) with violin plot displayed based on *INS* expression (log1(rpm+1)). **(B)** Barplot showing genes differentially expressed as log2FC and enriched in specific pathway from scβ-like-cells. **(C)** Scatter plot of 2670 hESC-derived *α*-like-cells from cluster 2 (**Fig. S4**) displayed based on *GCG* and *INS* expression (expo_logrpm). Total cell number from all genotypes combined. Encircled are monohormonal sc*α*-like-cells (Cluster 2a) (total of 641 cells) and bihormonal sc*α*-like-cells (Cluster 2b) (total of 1054 cells). **(D)** Violin plot of indicated cluster of cells displayed based on *GCG* and *INS* expression (log1(rpm+1)). Monohormonal scβ-like-cells (cluster 0) and bihormonal sc*α*-like-cells (cluster 2b) were identified and displayed as |logFC|>0.35 and adjusted p-value <1e^-4^. Monohormonal sc*α*-like-cells (cluster 2a) displayed as |logFC|>0.25 and adjusted p-value <1e^-2^. **(E)** Heatmap showing differentially expressed genes from monohormonal sc*α*-like-cells (Cluster 2a) by *HNF1A* genotypes. Total of 641 sc*α*-like-cells (cluster 2a) (all genotypes combined) were identified and displayed as |logFC|>0.25 and adjusted p-value <1e^-2^. (**F and G)** Barplot showing genes differentially expressed as log2FC and enriched in specific pathway from indicated endocrine cell type. **(H)** Heatmap showing top differentially expressed genes from bihormonal *α*/β-like-cells (Cluster 2b) across genotypes. Genes are listed in decreasing order of log_2_ fold change between *HNF1A* WT and *HNF1A* mutant genotypes. n=3 for each genotype. **(I-M)** Gene expression profile of *LINC01139* in scβ-like-cells, sc*α*-like-cells and cadaveric β-cells. **(I)** *LINC01139* gene expression levels (FPKM) determined by bulk RNAseq and after FACs sorting of *INS^GFP/wt^* positive cells from hESC-derived endocrine cells (hESC WT n=3, hESC Het n=3, hESC KO n=3 and MODY3 iPSC 460ins Het n=2). **(J)** *HNF1A* gene expression levels (FPKM) determined by single-cell RNAseq after FACs sorting of *INS^GFP/wt^* positive cells from hESC-derived endocrine cells (hESC WT n=113 cells and hESC KO n=158 cells). **(K)** Violin plot of scβ-like-cells and **(L)** sc*α*-like-cells based on *LINC01139* expression (log1(rpm+1)) by genotype. **(M)** *LINC01139* transcript levels of sorted pancreatic β-cells from non-diabetic donors (n=5) and MODY3 donor (n=1) (Haliyur et al. 2019). All stem cell differentiations were done for 27 days.

**Fig. S6.**
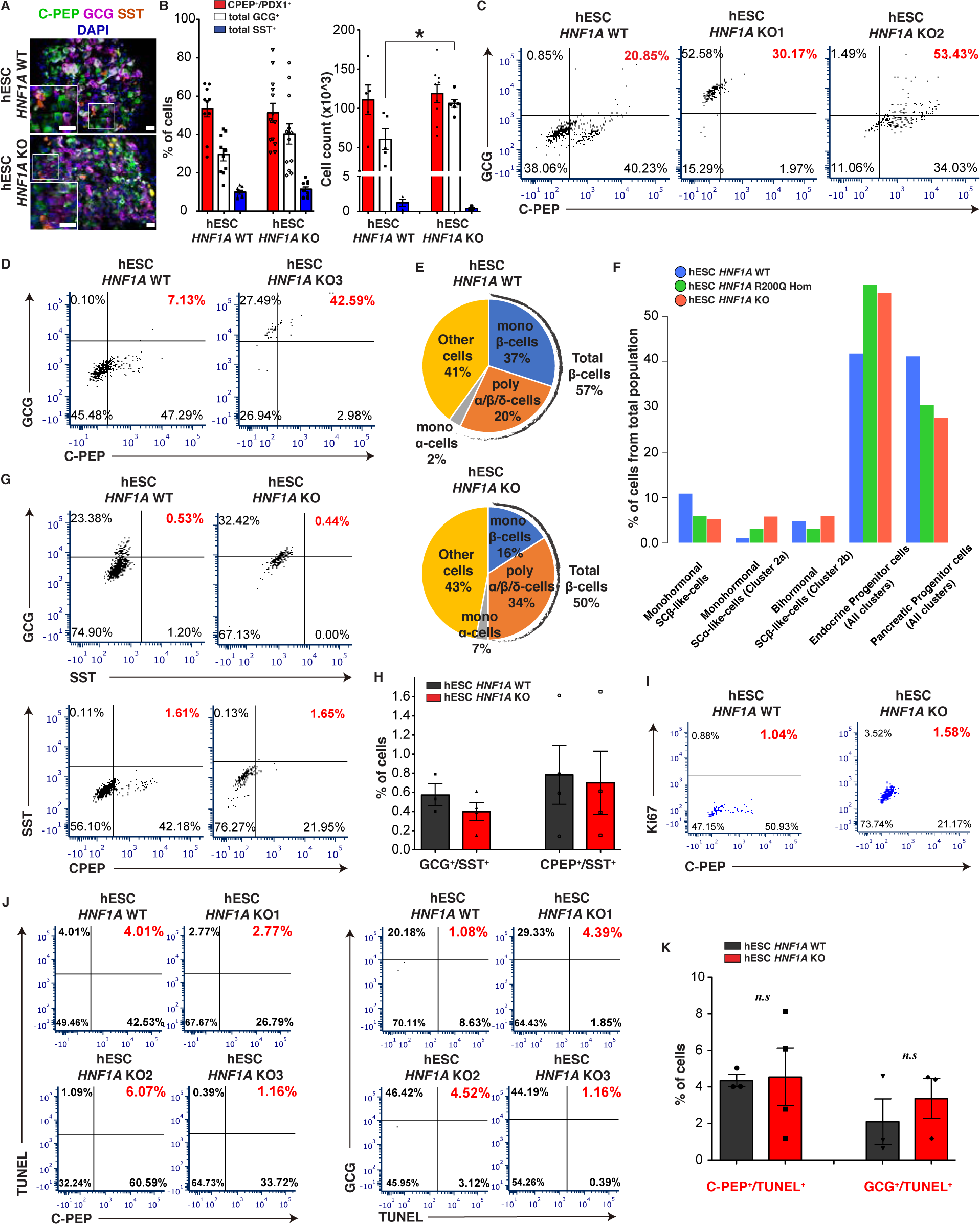
*HNF1A* deficiency causes a developmental bias towards the α-cell fate *in vitro.* **(A)** Representative IHC images of hESC-derived endocrine cell lines with indicated markers and **(B)** quantification of percentage and total cell count of scβ-like-cells (CPEP^+^/PDX1^+^ in red), sc*α*-like-cells (GCG^+^ in white) and scδ-like-cells (SST^+^ in blue). Cell count performed from 20 clusters. Scale bars: 20 µm. (**C and D)** Representative CPEP^+^ and GCG^+^ populations by flow cytometry from *HNF1A* KO1, KO2 and KO3. **(E)** Endocrine cells as percentage from total cells from hESC-derived clusters based on IHC staining. CPEP^+^/GCG^-^/SST^-^ cells indicate monohormonal β-cells and CPEP^+^/GCG^+^/SST^+^ cells indicate polyhormonal *α*/β/δ-cells. **(F)** Barplot showing the percentage (%) of indicated endocrine cell types from total cells (22164 cells) analyzed by single cell RNAseq from hESC-derived endocrine cells by *HNF1A* genotype (n=3 for each genotype). Endocrine progenitor cells are *SYP* expressing cells and pancreatic progenitor cells are *PDX1* expressing cells. **(G)** Representative GCG^+^, SST^+^ and CPEP^+^ populations by flow cytometry with **(H)** respective quantification. **(I and J)** Flow cytometry for indicated markers with **(K)** respective quantification. All stem cell differentiations were done for 27 days. 20 clusters (∼10k cells per cluster) of endocrine cells were used flow cytometry. For scatter plots, each point in plots represents an independent biological experiment (n). Data are represented as mean ± SEM. p-values: *p<0.05, **p<0.01, ***p<0.001; Mann-Whitney test. n.s: non-significant.

**Fig. S7.**
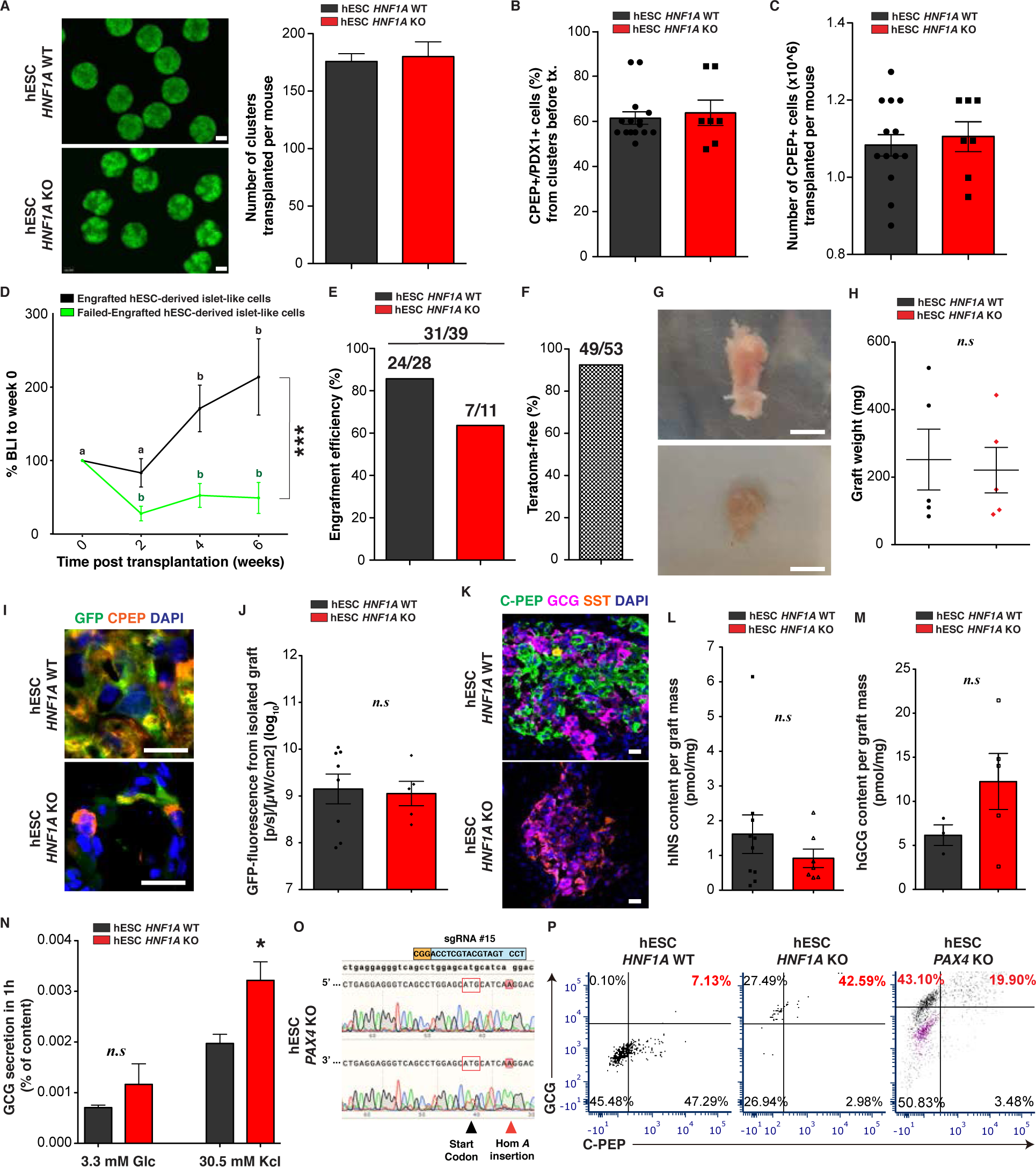
*HNF1A* deficiency causes a developmental bias towards the α-cell fate *in vivo*. **(A)** Representative image and number (WT n=17 and KO n=7) of pancreatic islet-like clusters transplanted per mouse and per genotype. Scale bars at 200 μm. **(B)** Percentage (%) of CPEP^+^/PDX1^+^ cells from clusters before transplantation with **(C)** total number of CPEP+ cells(10x6) transplanted per mouse and per genotype. **(D)** % Bioluminescence intensity (BLI) normalized to week 0 (transplantation day) of mice transplanted with *GAPDH^Luciferase/wt^* reporter hESC lines 0, 2, 4 and 6 weeks post-transplantation. Engrafted hESC-derived islet-like cells (WT n=11 and KO n=4) resulted in >50 pM of circulating human c-peptide after 6 weeks. Non-engrafted hESC-derived islet-like cells (WT n=4 and KO n=6) resulted in no or <5 pM of circulating human c-peptide after 30 weeks. Results are presented as mean of mice transplanted with both hESC genotypes. p-values were b: p<0.05. **(E)** Engraftment efficiency (%) from all mice transplanted with islet-like cells. **(F)** Teratoma-free mice transplanted with hESC-derived islet-like cell lines. Graft tissue was considered a teratoma when larger than 2 cm and by IHC of the germ layers. **(G)** Representative image of graft tissue 30 weeks post-transplantation with hESC-derived endocrine cells after explant from quadriceps muscle. Scale bars: 0.5 cm. **(H)** Graft weight in mg from explants. **(I)** Representative IHC image from isolated grafts for indicated markers. Scale bars: 20 µm. **(J)** GFP fluorescence ([p/s]/[µW/cm^2^)(log_10_) from isolated grafts. **(K)** IHC image from isolated graft for SCβ-cells (CPEP: c-peptide), sc*α*-cells (GCG: glucagon) and scδ-cells (SST: somatostatin). (**L and N**) Endocrine hormone content. **(L)** Human insulin and **(M)** human glucagon content per graft mass (pmol/mg) *ex vivo*. **(N)** Human glucagon secretion in 1h as % of content in static assay in response to indicated secretagogues from hESC-derived endocrine cells (WT n=4, KO n=7) *in vitro*. All protein concentrations were measured by ELISA**. (O)** Sanger sequencing results showing knockout frameshift mutation (A insertion) in hESC *PAX4* KO from both alleles in exon3 near start codon (red box). **(P)** Representative flow cytometry of CPEP^+^ and CGC^+^ populations. Gating for GCG and CPEP negative cells (magenta) was determined by incubating cells without primary antibodies and with secondary antibodies. 20 clusters (∼10k cells per cluster) of endocrine cells were used flow cytometry. All stem cell differentiations were done for 27 days for *in vitro* assay and for transplantation. All grafts were isolated 30 weeks post-transplantation for *ex vivo* analysis. For scatter plots, each point in plots represents an independent biological experiment (n). Data are represented as mean ± SEM. Different letters designate significant differences within group. p-values: *p<0.05, **p<0.01, ***p<0.001; Mann-Whitney test. n.s: non-significant.

**Fig. S8.**
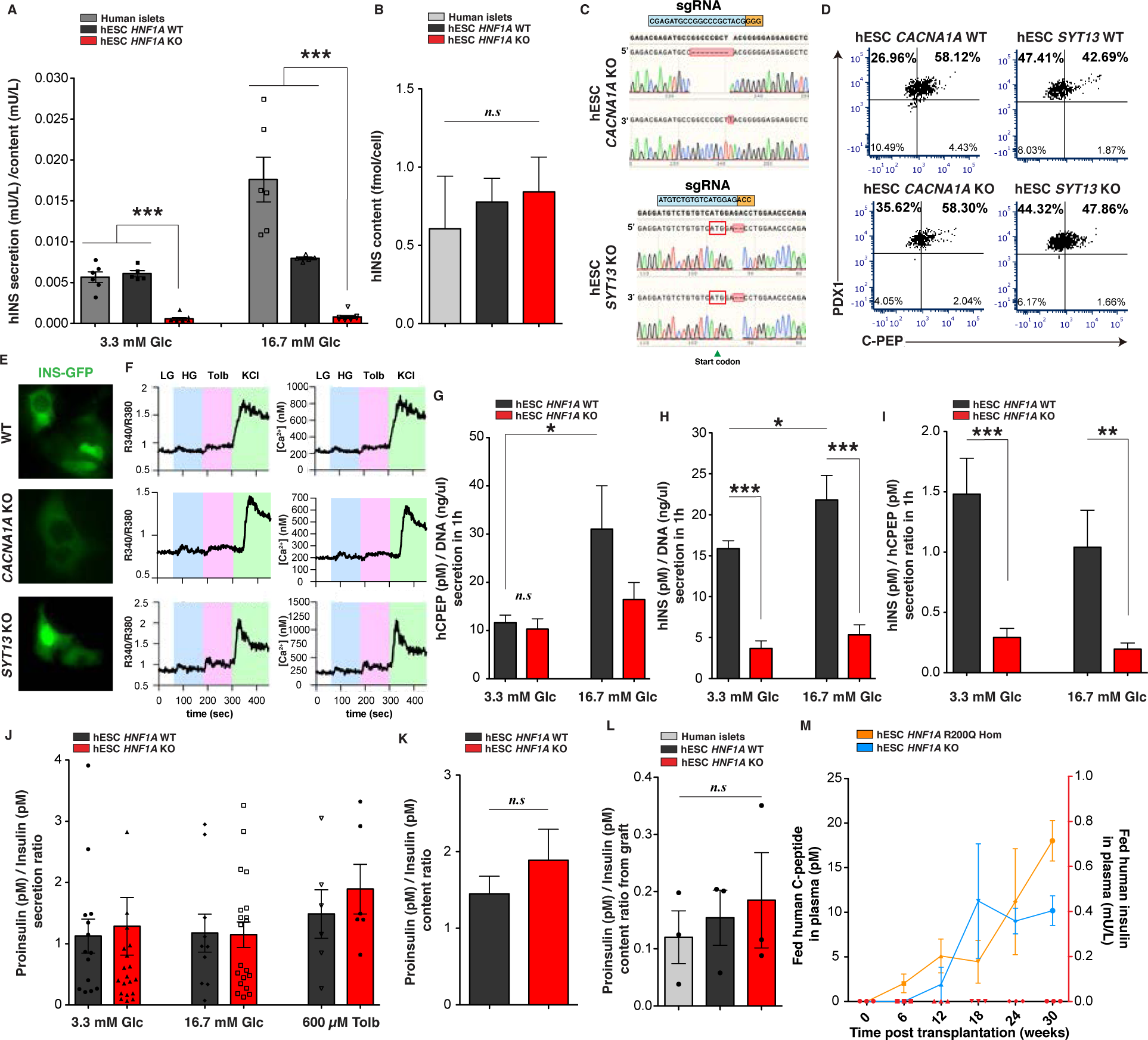
*HNF1A* deficiency affects insulin secretion in association with *CACNA1A* and *SYT13* down-regulation *in vitro*. **(A)** Human insulin secretion (mU/L) in 1h normalized to content (mU/L) in response to low glucose (LG, 3.3 mM) and high glucose (HG, 16.7 mM) stimulation in static assay. **(B)** Human insulin protein content per cell (fmol/cell) (Human islets n=6, WT n=17, KO n=21). **(C)** Sanger sequencing results showing knockout frameshift mutations in hESC *CACNA1A* KO and hESC *SYT13* KO line from both alleles near start codon (red box). **(D)** Representative flow cytometry of CPEP^+^ and PDX1^+^ populations. **(E)** dispersed scβ-like-cells (INS-GFP) with **(F)** representative intracellular calcium levels measured as R340/R380 and [Ca^2+^](nM) over time (sec) with indicated secretagogues. **(G)** Human c-peptide (pM) and **(H)** human insulin secretion (pM) in 1h normalized to DNA (ng/µl) in response to basal (3.3 mM) and high (16.7 mM) glucose stimulation in static assay (WT n= 7 and KO n=20). **(I)** hINS(pM)/hCPEP(pM) secretion ratio in response to basal (3.3 mM) and high (16.7 mM) glucose stimulation (WT n=9 and KO n=14). **(J-L)** Proinsulin(pM)/Insulin(pM) ratio from **(J)** secretion and **(K)** content from *in vitro* cells (WT n=8 and KO n=15) and **(L)** *ex vivo* grafts. (**M**) Plasma human C-peptide (pM) (left y-axis) and human insulin secretion (mU/L) (right y-axis) in plasma of *ad libitum-fed* mice transplanted with hESC *HNF1A* KO1 and *HNF1A* R200Q homozygous-derived endocrine cells (n=3). All stem cell differentiations were done for 27 days. All protein concentrations were measured by ELISA. For scatter plots, each point in plots represents an independent biological experiment (n). Data are represented as mean ± SEM. p-values: *p<0.05, **p<0.01, ***p<0.001; Mann-Whitney test. n.s: non-significant.

**Fig. S9.**
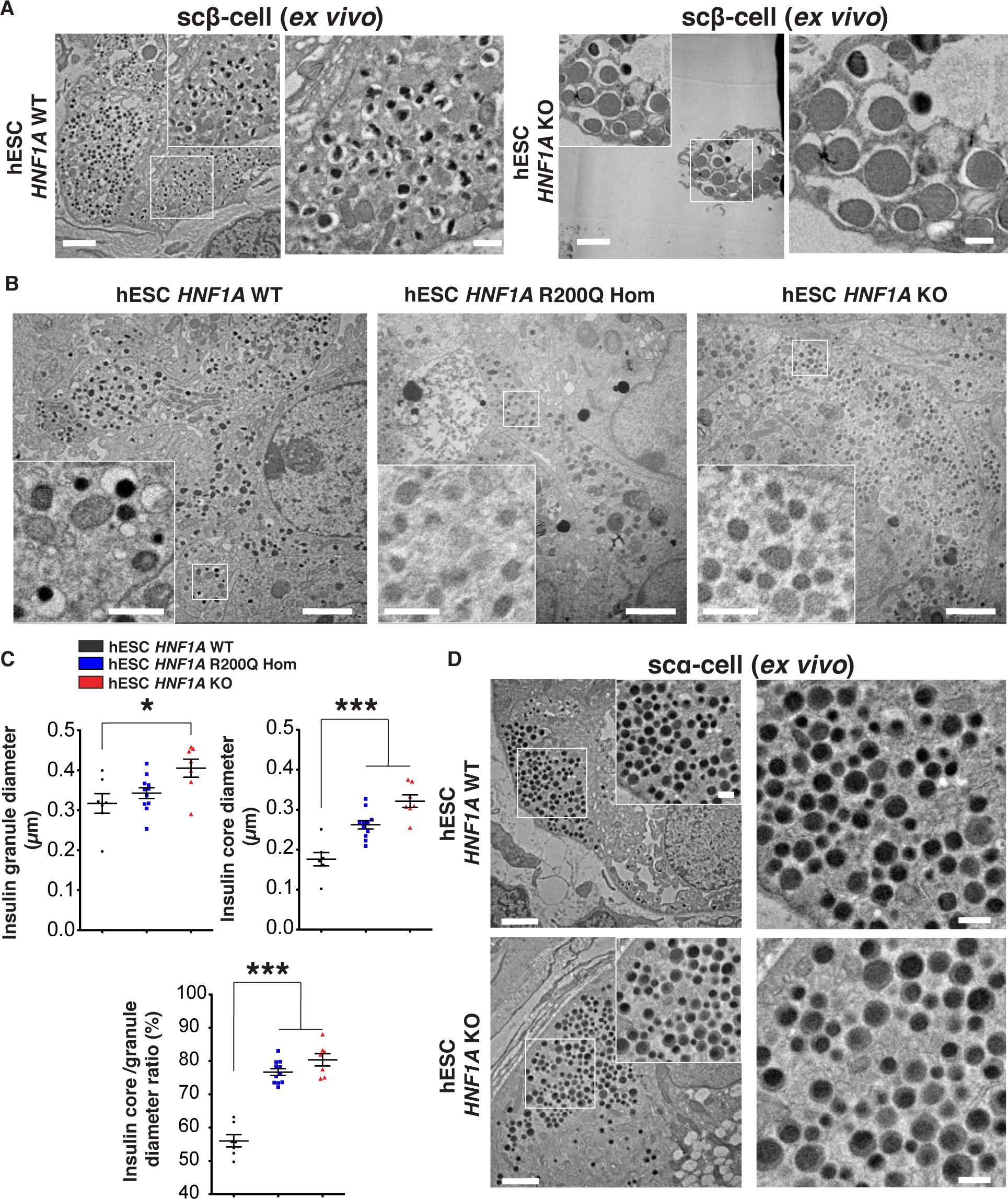
*HNF1A* deficiency causes abnormal insulin granule structure. (**A**) Representative electron microcopy (EMC) image of scβ-cells *ex vivo* from isolated graft 30 weeks post-transplantation. Explants are from euglycemic mice. **(B)** Representative EMC images of scβ-like-cells *in vitro* with **(C)** quantification of insulin granule diameter (µm), insulin granule core diameter (µm) and insulin granule core diameter to insulin granule diameter ratio (%) per cell. **(D)** Representative EMC images of sc*α*-cells. For all images, scale bars: 2 µm in low and 0.5 µm in high magnification. Each point in plots is the average of insulin granules per scβ-like-cells. All stem cell differentiations were done for 27 days for *in vitro* assay and for transplantation. For scatter plots, each point in plots represents an independent biological experiment (n). Data are represented as mean ± SEM. p-value: *p<0.05, **p<0.01, ***p<0.001; two-tailed t-test. n.s: non-significant.

**Fig. S10.**
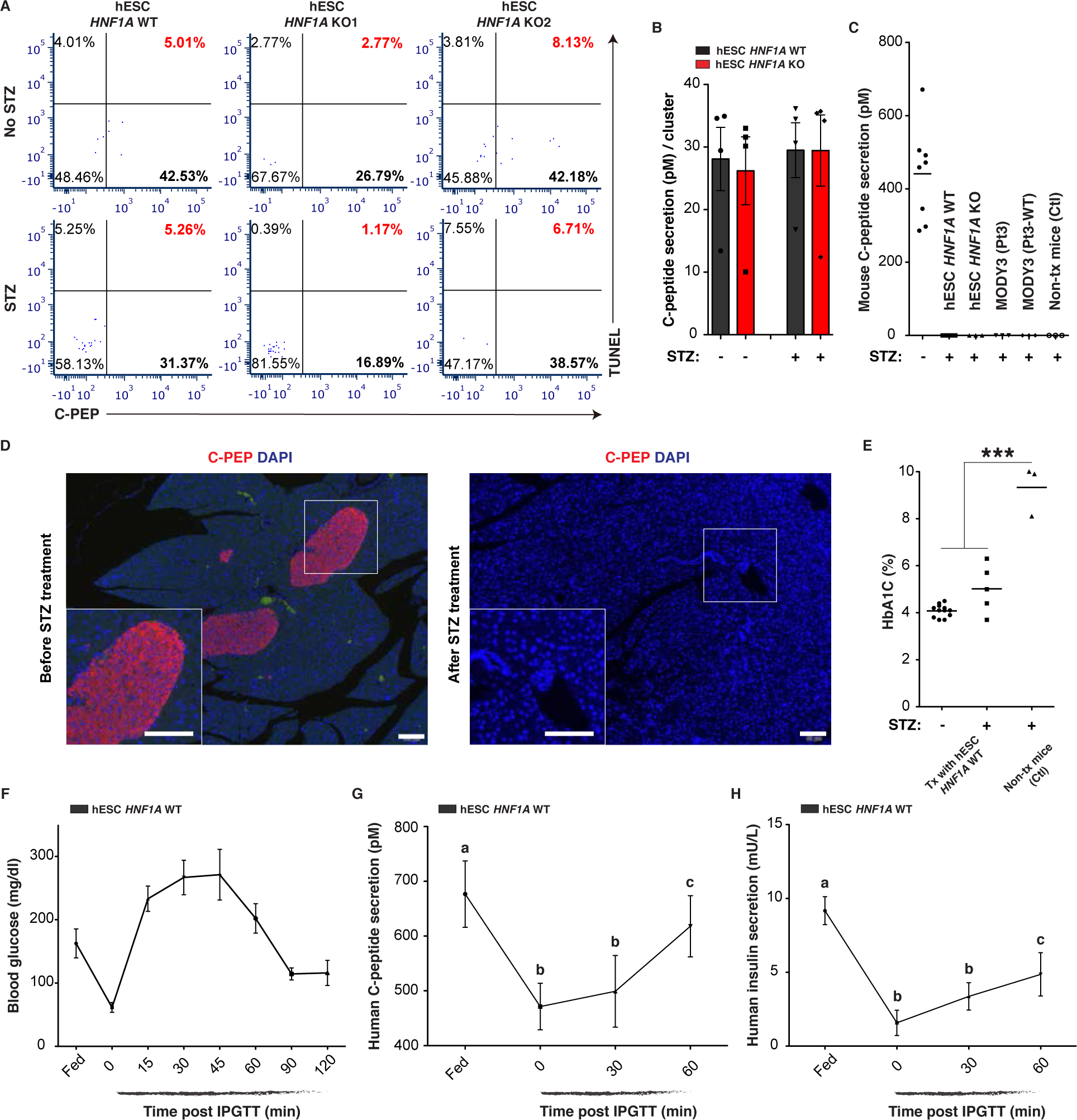
*HNF1A* deficient scβ-cells are unable to maintain glucose homeostasis in diabetic mice. **(A and B)** hESC-derived endocrine cells were treated with (4mg/ml; 15mM) or without STZ *in vitro*. **(A)** Apoptosis quantification after flow cytometry of TUNEL^+^ and CPEP^+^ cells. 10 clusters (∼10k cells per cluster) of endocrine cells were used flow cytometry. **(B)** Human c-peptide secretion (pM) per cluster measured before and after STZ *in vitro*. **(C)** Mouse C-peptide secretion (pM) in plasma of *ad libitum-fed* mice transplanted with indicated cell lines or non-transplanted mice (Ctl) before (-) or >3 weeks after STZ injection (+). **(D)** Representative immunofluorescent staining of mouse pancreas showing islets (CPEP; red) before and after STZ treatment. Scale bars: 20 µm. **(E)** HbA1C (%) in mice transplanted with hESC *HNF1A* WT-derived endocrine cells or without cells (Ctl) before (-) and 5-15 weeks after (+) STZ treatment. (**F-H**) IPGTT in mice transplanted with hESC *HNF1A* WT-derived endocrine cells (n=6) several weeks (>3 weeks) after STZ treatment in *ad libitum-fed* state and during an iPGTT (t0, t30 and t60). **(F)** Blood glucose concentrations (mg/dl), **(G)** human C-peptide (pM) and **(H)** human insulin secretion (mU/L) in plasma. p-values were b: p<0.001, c: p<0.05. All stem cell differentiations were done for 27 days. For scatter plots, each point in plots represents an independent biological experiment (n). Data are represented as mean ± SEM. Different letters designate significant differences within group. p-values: *p<0.05, **p<0.01, ***p<0.001; two-tailed t-test.

**Fig. S11.**
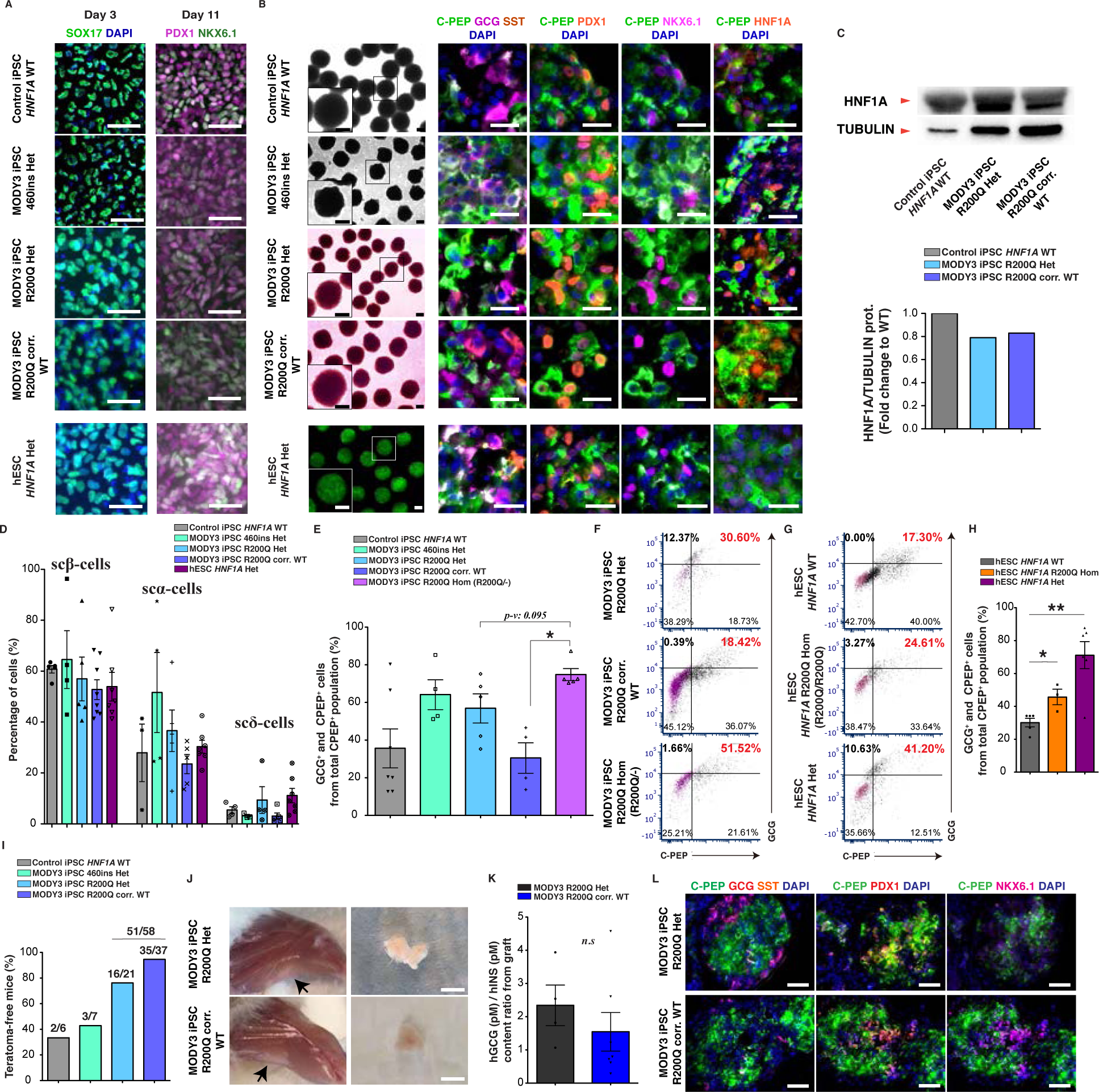
*HNF1A* R200Q mutation is pathogenic and causes developmental bias towards the *α*-cell fate *in vitro*. (A) Representative IHC staining of stem cell-derived cell lines *in vitro* at definitive endoderm stage with SOX17 (day 3), pancreatic progenitor stage with PDX1 and NKX6.1 (day 11) and endocrine cell stage (day 27). Scale bars: 50 µm. White cells are PDX1/NKX6.1 double positive cells. (B) Representative bright-field images from MODY3 iPSC-derived organoids and GFP image from hESC Het-derived organoids with IHC staining for indicated markers. White cells are GCG/CPEP double positive cells. Scale bars: 20 µm. (C) Western blot for HNF1A (67 kDa) and β-Tubulin-III (55 kDa) from iPSC-derived endocrine cells with respective quantification on day 27 of differentiation *in vitro*. (D) Percentage of scβ-like-cells (CPEP^+^/PDX1^+^), sc*α*-like-cells (GCG^+^) and scδ-like-cells (SST^+^) determined by IHC. (E) Quantification of GCG and CPEP positive cells from total CPEP positive cell population determined by IHC. (F) CPEP^+^ and GCG ^+^ populations from iPSC-derived endocrine cells and (G) hESC-derived endocrine ells by flow cytometry. Gating for GCG and CPEP negative cells (magenta) was determined by incubating cells without primary antibodies and with secondary antibodies. (H) Quantification of GCG and CPEP positive cells from total CPEP positive cell population (%) determined by flow cytometry. (I) Teratoma-free mice transplanted with MODY3 iPSC-derived endocrine cells. Graft tissue was considered a teratoma when larger than 2 cm and by IHC of the germ layers. (J) Representative image of graft tissue (black arrows) 30 weeks post-transplantation with MODY3 iPSC-derived endocrine cells before and after explant from quadriceps muscle. Scale bars: 0.5 cm. (K) Human glucagon content per graft mass (pmol/mg) *ex vivo* measured by ELISA. (L) Representative IHC images showing MODY3 iPSC-derived endocrine cells from isolated graft. Scale: 50 µm. All stem cell differentiations were done for 27 days for *in vitro* assay and for transplantation. All grafts were isolated 30 weeks post-transplantation for *ex vivo* analysis. For scatter plots, each point in plots represents an independent biological experiment (n). Data are represented as mean ± SEM. Different letters designate significant differences within group. p-values: *p<0.05, **p<0.01, ***p<0.001; Mann-Whitney test. n.s: non-significant.

**Fig. S12.**
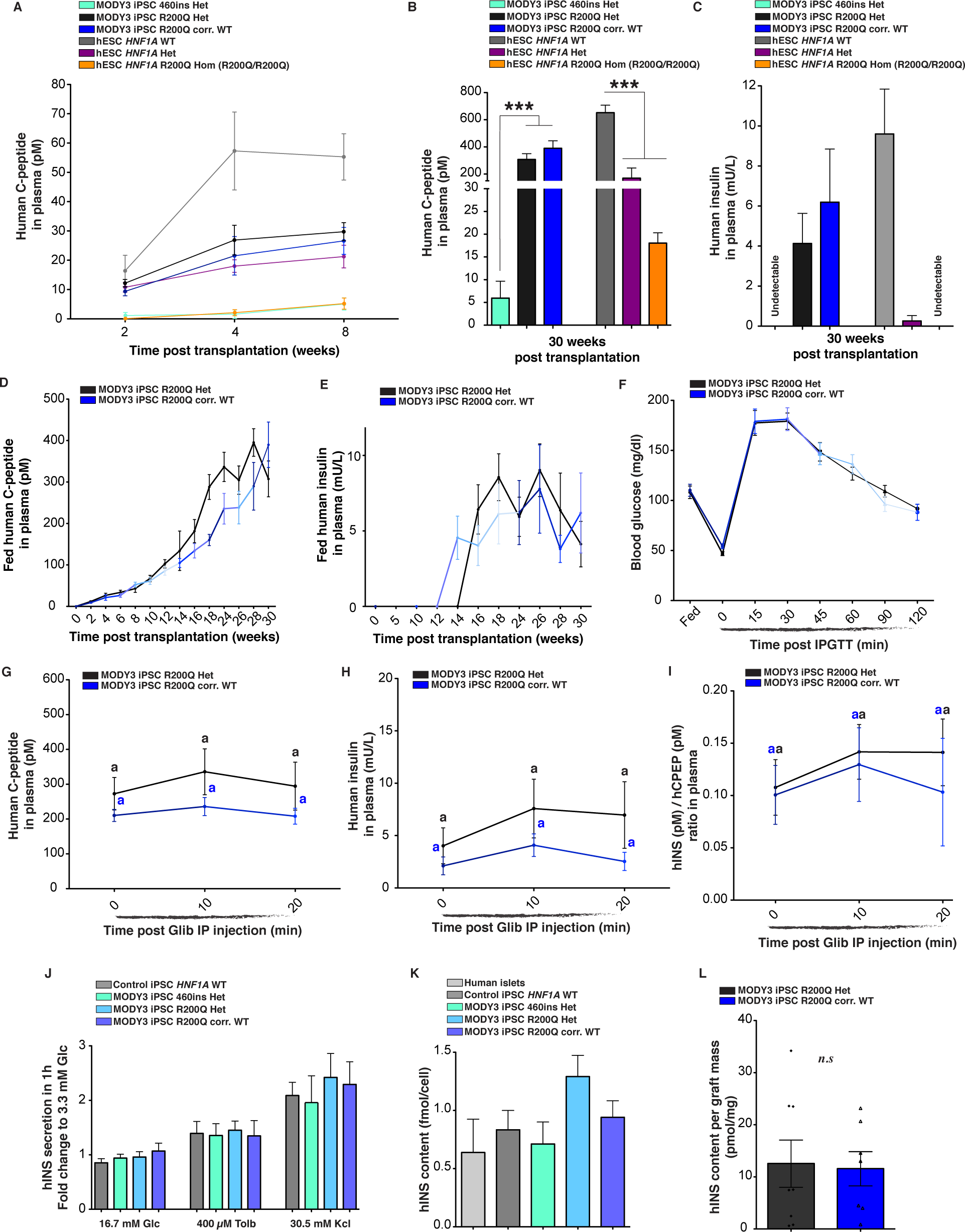
MODY3 iPSC-derived β-cells are initially glucose responsive. (**A-C**) Human C-peptide (pM) and human insulin levels in plasma of *ad libitum-fed* mice transplanted with MODY3 iPSC- and hESC-derived endocrine cells**. (A)** Plasma human C-peptide (pM) two, four and eight weeks post transplantation in mice transplanted with MODY3 iPSC lines (460ins Het n=4, R200Q Het n=12 and R200Q corr. WT n=12) and hESC lines (WT n=17, Het n=10 and R200Q Hom n=6). **(B)** Plasma human C-peptide (pM) thirty weeks post transplantation in mice transplanted with MODY3 iPSC lines (460ins Het n= 5, R200Q Het n=5 and R200Q corr. WT n=6) and hESC lines (WT n=7, Het n=4 and R200Q Hom n=3) and **(C)** human insulin (mU/L) thirty weeks post transplantation in mice transplanted with MODY3 iPSC lines (460ins Het n= 3, R200Q Het n=4 and R200Q corr. WT n=6) and hESC lines (WT n=8, Het n=7 and R200Q Hom n=3). **(D)** Plasma human C-peptide (pM) and **(E)** human insulin secretion (mU/L) in plasma of *ad libitum-fed* mice transplanted with MODY3 iPSC-derived endocrine cells (R200Q Het n=14 and R200Q corr. WT n=12). **(F)** Blood glucose concentrations (mg/dl) in *ad libitum-fed* state and during an iPGTT (t0, t30 and t60) in mice with MODY3 iPSC-derived endocrine cells (R200Q Het n=12 and R200Q corr. WT n=10). **(G-I**) Glibenclamide injection in *ad libitum-fed* mice transplanted with MODY3 iPSC-derived endocrine cells (R200Q Het n=5 and R200Q corr. WT n=9). **(G)** Human C-peptide secretion (pM), **(H)** human insulin secretion (mU/L) and **(I)** hINS (pM)/hCPEP (pM) secretion ratios. **(J)** Human insulin secretion in 1h from static assay as a fold change to 3.3 mM Glc in response to indicated secretagogues *in vitro* (iPSC WT n=5, 460ins Het n=5, R200Q Het n=10 and R200Q corr. WT n=7). **(K)** Human insulin protein content per cell (fmol/cell) *in vitro* (Human islets n=6, iPSC WT n=4, 460ins Het n=4, R200Q Het n=7 and R200Q corr. WT n=5). **(L)** Human insulin content per graft mass (pmol/mg). All stem cell differentiations were done for 27 days for *in vitro* assay and for transplantation. All grafts were isolated 30 weeks post-transplantation for *ex vivo* analysis. All protein concentrations were measured by ELISA. For scatter plots, each point in plots represents an independent biological experiment (n). Data are represented as mean ± SEM. Different letters designate significant differences within group. p-values: *p<0.05, **p<0.01, ***p<0.001; two-tailed t-test. n.s: non-significant.

**Fig. S13.**
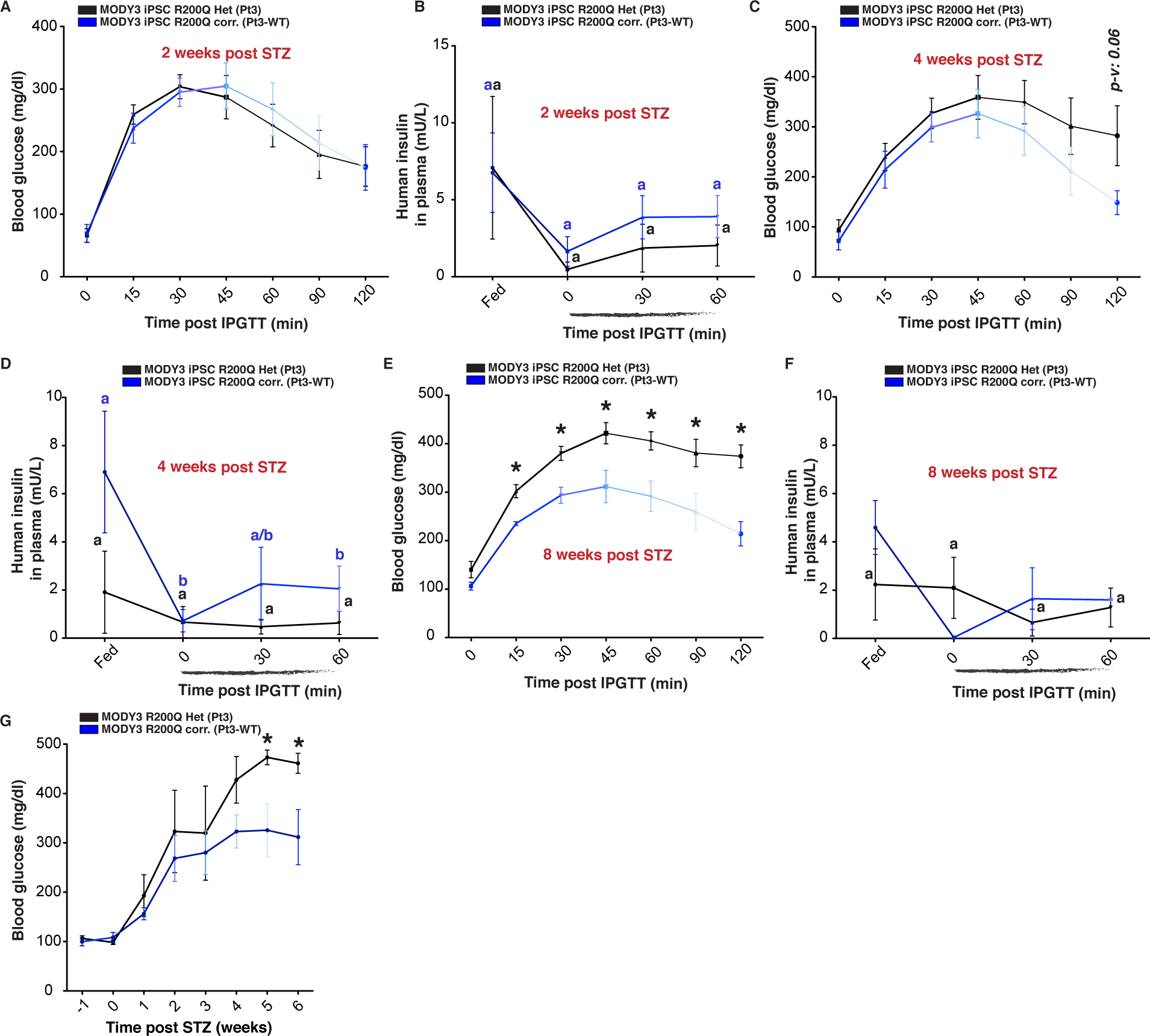
*HNF1A* haploinsufficiency gradually impairs sc*β*-cell function *in vivo.* (**A-F**) IPGTT in mice transplanted with MODY3 iPSC-derived endocrine cells in *ad libitum-fed* state and during an iPGTT (t0, t30 and t60) 2, 4 and 8 weeks post STZ treatment. **(A)** Blood glucose concentrations (mg/dl) and **(B)** human insulin secretion (mU/L) in plasma of mice after 2 weeks post STZ (R200Q Het n=12; R200Q corr. WT n=10). **(C)** Blood glucose concentrations (mg/dl) and **(D)** human insulin secretion (mU/L) in plasma of mice after 4 weeks post STZ (R200Q Het n=5; R200Q corr. WT n=4). **(E)** Blood glucose concentrations (mg/dl) and **(F)** human insulin secretion (mU/L) in plasma of mice after 8 weeks post STZ (R200Q Het n=5; R200Q corr. WT n=2). p-values were b and c: p<0.05. **(G)** Blood glucose levels (mg/dl) monitored in *ad libitum-fed* mice weeks before (-1) or after (0 to 6) STZ injection (R200Q Het n=10; R200Q corr. Wt n=7). All protein concentrations were measured by ELISA. For scatter plots, each point in plots represents an independent biological experiment (n). Data are represented as mean ± SEM. Different letters designate significant differences between fed, t0, t30 and t60 for each genotype. p-values: *p<0.05,**p<0.01, ***p<0.001; two-tailed t-test.

